# Activation of alternative oxidase ensures carbon supply for ethylene and carotenoid biosynthesis during tomato fruit ripening

**DOI:** 10.1101/2025.01.30.635696

**Authors:** Ariadna Iglesias-Sanchez, Nestor Fernandez Del-Saz, Miguel Ezquerro, Elisenda Feixes-Prats, Miquel Ribas-Carbo, Alisdair R. Fernie, Manuel Rodriguez-Concepción, Igor Florez-Sarasa

## Abstract

Tomato (*Solanum lycopersicum*) is a climacteric fruit displaying a peak of respiration at the onset of ripening accompanied by increased synthesis of ethylene and carotenoid pigments. Chromoplast and mitochondrial respiration participate at different stages of fruit ripening, but their *in vivo* regulation and function remains unclear. We determined the *in vivo* activities of the mitochondrial alternative oxidase (AOX) and cytochrome oxidase pathways and quantified the levels of respiratory- and ripening-related gene transcripts, primary metabolites and carotenoids in ripening tomato fruits with or without a functional chromorespiration. Furthermore, we carried out physiological, molecular and metabolic analyses of CRISPR-Cas9 mutants defective in AOX1a, the main AOX isoform up-regulated during tomato fruit ripening. We confirmed that PTOX-dependent chromorespiration is only relevant at late stages of ripening and found that *in vivo* AOX activity significantly increased at the breaker stage, becoming the main contributor to climacteric respiration when ripening is unleashed. This activation did not correlate with gene expression but was likely due to increased levels of AOX activators such as pyruvate (a metabolic precursor of carotenoids), 2-oxoglutarate and succinate. A strong alteration of ripening-related metabolites was observed in *aox1a* mutant fruits, highlighting a key role of the AOX pathway at the onset of ripening. Our data suggest that increased supply of TCA cycle intermediates at climacteric stage allosterically enhance AOX activity, thus allowing the reoxidation of NAD(P)H to ensure carbon supply for triggering ethylene and carotenoid biosynthesis.

## INTRODUCTION

Tomato (*Solanum lycopersicum*) is the main model system for fleshy fruit ripening studies at biochemical, genetic and molecular levels (Giovannoni et al., 2017; Zhu et al., 2018; Li et al., 2019; Kaur et al., 2023). In this economically relevant fruit, a sudden increase in respiration occurs at the onset of ripening, usually in concert with increased ethylene production that eventually impacts fruit color, firmness, taste, and flavor (Klee and Giovannoni, 2011). Tomato ripening involves the degradation of chlorophylls present in mature green fruit and the accumulation of high levels of carotenoid pigments. As a result, chloroplasts differentiate into specialized plastids named chromoplasts and the fruit color changes from green to red when ripe (Giovannoni et al., 2017). The supply of energy (ATP) and carbon precursors for the synthesis of carotenoids and other health-promoting metabolites during fruit ripening is assumed to rely on sugar import from source tissues. Chromorespiration, a respiratory process located in chromoplasts that involves the NAD(P)H dehydrogenase complex and the plastid terminal oxidase (PTOX), was shown to provide ATP in tomato fruits at late ripening stages (Renato et al., 2014). However, respiratory pathways acting in the cytosol and mitochondria are proposed to be the main suppliers of ATP and metabolic precursors at earlier stages (Bouvier and Camera, 2007; Beauvoit et al., 2018; Pott et al., 2019).

In plants, respiration involves three main steps: glycolysis in the cytosol, the tricarboxylic acid (TCA) cycle in the mitochondrial matrix, and the electron transport chain in the inner mitochondrial membrane (Fernie et al., 2004). The last step includes alternative respiratory pathways involving internal and external type II NAD(P)H dehydrogenases, uncoupling proteins (UCPs) and alternative oxidases (AOXs) (Moller et al., 2021). AOX constitutes the cyanide-insensitive pathway that forks away from the cytochrome c oxidase (COX) pathway at the level of ubiquinone (UQ) bypassing proton pumping through complexes III and IV (Del-Saz et al., 2018; Moller et al., 2021), whereas UCPs uncouple the proton gradient(Vercesi et al., 2006; Barreto et al., 2020). Both the redox and the proton electrochemical gradient energy-dissipating pathways (AOX and UCP, respectively) lead to a decrease in ATP synthesis. However, AOX and UCP activities are beneficial for plant performance under different physiological conditions mainly due to their roles in providing metabolic flexibility and in lowering the level of mitochondrial reactive oxygen species (ROS) (Del-Saz et al., 2018; Barreto et al., 2020). Several AOX and UCP homologs have been reported in tomato (Holtzapffel et al., 2003; Vercesi et al., 2006; Xu et al., 2012) and some of them have been proposed to participate in tomato fruit ripening (Almeida et al., 1999; Holtzapffel et al., 2003; Xu et al., 2012). However, the precise contribution of the different AOX isoforms during climacteric respiration as well as their role in fruit metabolism (including the production of carotenoid pigments) remains poorly understood.

In tomato, there are four AOX-encoding genes: three *AOX1* (*AOX1a*, *AOX1b*, and *AOX1c*) and one *AOX2*. From them, only *AOX1a* and, to a lower extent, *AOX1b* were reported to be up-regulated during tomato fruit ripening (Xu et al., 2012). Although the selective expression of AOX isoforms can influence its overall activity, a crucial layer of AOX regulation involves posttranslational modifications which mainly involve changes in redox status and allosteric interactions with organic acids (Selinski et al., 2018; Florez-Sarasa et al., 2019). Notably, *in vitro* studies have shown that the activation of AOX by different organic acids from the TCA cycle is isoform-specific (Selinski et al., 2018). An additional challenge for understanding the role of AOX in fruit ripening is the lack of fundamental information about the *in vivo* electron partitioning between the AOX and the COX pathways during fruit ripening. Early studies assessed tomato fruit respiration by using specific respiratory inhibitors (Theologis and Laties, 1978; Xu et al., 2012), which are not adequate for measuring *in vivo* respiratory activities (Day et al., 1996; Del-Saz et al., 2018). Alternatively, *in vivo* electron partitioning between the AOX and the COX pathways can be determined using the O_2_ isotope (^18^O) discrimination technique (Del-Saz et al., 2017, 2018), which allows for precise estimations of the energy-efficiency of respiration. The ^18^O discrimination technique has been applied to study respiration in different species and tissues exposed to many different conditions (Del-Saz et al., 2017, 2018). However, the application of this technique in fruit tissues has been limited due to (i) the limitation of O_2_ diffusion due to pericarp thickness (Del-Saz et al., 2017) and (ii) the interference of non-mitochondrial O_2_ consumption, mainly due to wounding respiration (Theologis and Laties, 1978; Moller et al., 1988) and chromorespiration (Renato et al., 2014). These and other technical difficulties in measuring fruit respiration have been recognized and bypassed by using a stoichiometric model, which predicted a peak of respiratory flux (CO_2_ released) and energy dissipation just before ripening and coinciding with the climacteric peak of respiration (Colombié et al., 2017). While this model has been used to calculate theoretical AOX- and UCP-associated respiratory fluxes during fruit ripening in different conditions (Colombié et al., 2017), experimental confirmation is missing.

In the present study, we have used wild-type tomato plants (cv. Ailsa Craig) as well as PTOX-defective *ghost* mutants with no chromorespiration (Barr et al., 2004) to experimentally determine the *in vivo* activities of the AOX and COX pathways during tomato fruit ripening. These data in combination with transcript and metabolite profiling revealed key regulations of the *in vivo* AOX activity at the climacteric stage. Furthermore, intensive respiration, transcript and metabolomics analysis throughout several ripening stages were performed in newly generated *aox1a* knockout mutants to study the role of the AOX pathway during fruit ripening. Our results strongly suggest a crucial role of the AOX pathway on the provision of carbon skeletons during fruit ripening, particularly for ethylene synthesis.

## RESULTS

### Measurements of oxygen consumption and isotope discrimination during tomato fruit ripening

Respiration under our experimental plant growth conditions was initially assessed with an oxygen electrode in sliced pericarp tissues from wild-type (WT) Ailsa Craig tomato fruits at four different stages of development selected based on color: mature green (MG), breaker (BR), orange (OR) and red (R) (Fig. 1A). Oxygen consumption was determined after different times of incubation with the respiration buffer containing CaCl_2_ to avoid and wash out the so called ‘wounding’ or ‘residual respiration’ (Theologis and Laties, 1978; Moller et al., 1988). Respiration rates among different time incubation points were not statistically different (P<0.05) at any developmental stage (Supplemental Fig. S1A). An incubation time of 20 min prior to the measurement of respiration was chosen for all subsequent respiration analyses. To next investigate the contribution of different respiratory pathways, we tested the effects of respiratory inhibitors on oxygen consumption rates (Supplemental Fig. S1B). Octyl-gallate (OGAL) at 1 mM concentration was reported to inhibit O_2_ consumption in tomato fruit pericarp only at late stages of ripening (OR and R), likely due to a specific effect on plastid terminal oxidase (PTOX) and not on AOX (Renato et al., 2014). However, our experiments using 1 mM OGAL in the presence of the COX inhibitor potassium cyanide (KCN) do not support this conclusion (Supplemental Fig. S1B). The KCN-resistant respiration at MG and BR stages (which lack PTOX activity) was substantially reduced after OGAL treatment, thus suggesting a strong inhibitory effect of OGAL on AOX (Supplemental Fig. S1B). Consistent with this conclusion, adding the AOX inhibitor salicylhydroxamic acid (SHAM) did not alter the effect of KCN plus OGAL treatments in MG and BR fruit samples (Supplemental Fig. S1B).

**Fig. 1.**
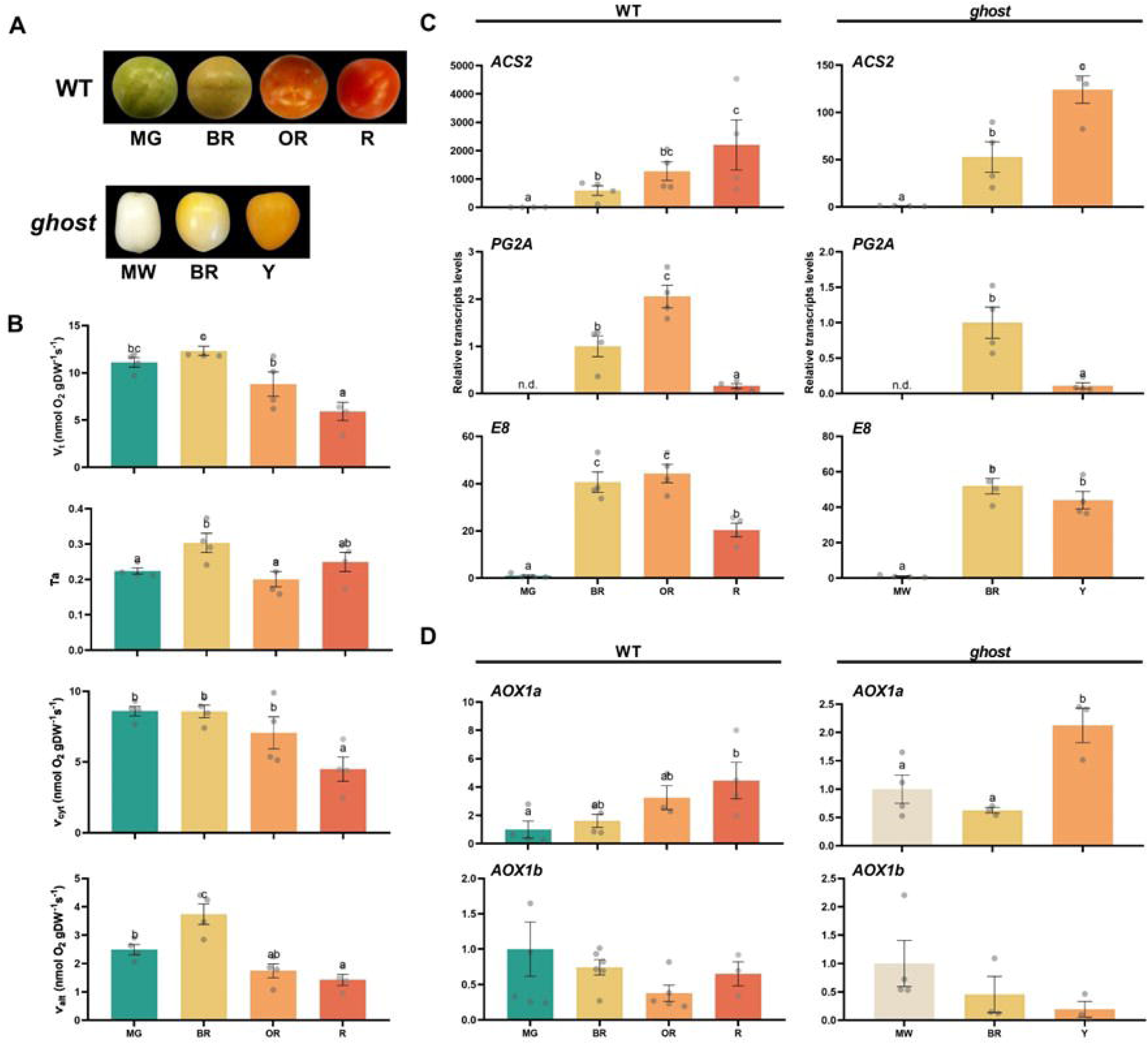
Visual phenotypes, *in vivo* respiration and gene expression in wild-type (WT, cv Ailsa Craig) and *ghost* mutant. (A) WT and *ghost* tomato fruits at different ripening stages: MG, mature green; BR, breaker; OR, orange; R, red; MW, mature white; Y, yellow. (B) Total respiration (V_t_), electron partitioning to the alternative oxidase pathway (τ_a_), cytochrome oxidase pathway activity (ν_cyt_), and alternative oxidase pathway activity (ν_alt_). Relative transcript levels of (C) ripening-related genes (E8: 2-oxoglutarate-dependent dioxygenase ethylene-responsive protein; ACS2: aminocyclopropane-1-carboxylate synthase 2; PG2a: polygalacturonase 2a) and (D) AOX-related genes (AOX1a: alternative oxidase 1a; AOX1b: alternative oxidase 1b) in WT and *ghost* mutant fruits at different ripening stages. Data are presented as fold-changes relative to MG or MW stages in WT and *ghost* fruits, respectively (i.e., levels at MG and MW stages are set to 1). All values are means ± SE of 3 to 6 replicates. Significant differences (P < 0.05) are indicated by different letters. “n.d.” denotes “not detected.“

Because respiratory inhibitors do not allow to properly measuring *in vivo* respiratory activities (Day et al., 1996; Del-Saz et al., 2018), we used the O_2_ isotope (^18^O) discrimination technique (Del-Saz et al., 2017, 2018) to investigate the *in vivo* electron partitioning between the AOX and the COX pathways. To measure ^18^O discrimination in fruit pericarp tissue, we initially optimized sample preparation in the absence (Δ_n_) and presence of the respiratory inhibitors KCN (Δ_a_, ^18^O discrimination of AOX) and SHAM (Δ_c_, ^18^O discrimination of COX). When pericarp samples were sliced into pieces of different thickness, results were variable and values were lower than those previously published for several tissues and species, i.e. Δ_c_ and Δ_n_ between 18 and 20 ‰ and Δ_a_ between 26 and 30 ‰ (Ribas-Carbo et al., 2005) (Supplemental Fig. S2). This might be caused to diffusion through dense or thick tissues (Del-Saz et al., 2017). Following previous procedures with other thick tissues (Gomez-Casanovas et al., 2007), we sliced the fruit pericarp samples in ca. 2 mm thick and 1 cm long pieces (Supplemental Fig. S2). This resulted in ^18^O discrimination values that were less variable and, most importantly, within the expected range (Supplemental Fig. S2).

Based on the described results, we next carried out the assessment of ^18^O discrimination in the absence and presence of respiratory inhibitors at different fruit ripening stages (Table 1). The ^18^O discrimination in the presence of OGAL (Δ_c-OGAL_) at the MG stage was lower than in the absence of inhibitors (Δ_n_) but similar to that SHAM-treated (Δ_c-SHAM_) samples (Table 1), thus demonstrating that both SHAM and OGAL had a similar inhibitory effect on AOX, in agreement with our previous data (Supplemental Fig. S1B). Furthermore, Δ_c-OGAL_ values in the range of those typically reported as end-point COX values (i.e. 18-20 ‰) were found at all ripening stages, thus confirming that OGAL inhibits *in vivo* AOX activity and hence should not be used to specifically inhibit only PTOX.

**Table 1.**
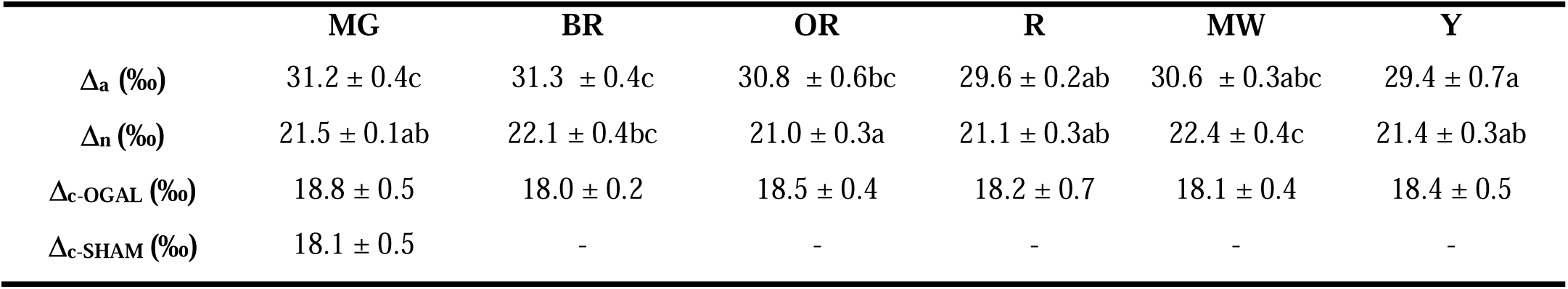
Oxygen isotope discrimination (‰) in the absence of inhibitors (Δ_n_), in the presence of 20 mM SHAM (Δ_c-SHAM_), 5 mM KCN (Δ_a_) and 1 mM OGAL (Δ_c-OGAL_) in mature green (MG), breaker (BR), orange (OR) and red (R) fruits from wild-type tomato plants as well as in mature white (MW) and yellow ripened (Y) fruits from ghost plants. Values are mean ± SE of three to six replicates. Significant (p < 0.05) differences between species are denoted by different letters.

As a complementary strategy to evaluate the contribution of chromorespiration to the total respiration of the tomato fruit pericarp at different ripening stages, we used the tomato PTOX-defective *ghost* mutant (Barr et al., 2004). Because of the absence of chlorophylls and carotenoid pigments in *ghost* fruits, we refer to their fruit development stages as mature white (MW) when reached full size, breaker (BR) when yellow flavonoid pigments start to accumulate, and yellow (Y) when fruit becomes fully colored (Fig. 1A). Δ_c-OGAL_ was very similar in WT and *ghost* tomatoes at all ripening stages, thus reinforcing our previous conclusion on the inhibitory effect of OGAL on AOX. Notably, Δ_a_ was also similar in WT and *ghost* fruits at both non-ripe (MG and MW) and ripe (R and Y) stages (Table 1). Therefore, these results suggest a minor contribution of PTOX-based chromorespiration to O_2_ consumption in the fruit pericarp during ripening.

Next, ^18^O discrimination data (Table 1) were used to determine the electron partitioning to AOX (τ_a_) in WT fruit at each ripening stage (Fig. 1B). There was a significant (P<0.05) increase in τ_a_ at the BR stage as compared to the MG stage (Fig. 1B). Total respiration rate (V_t_) also increased at BR stage, but it significantly (P<0.05) decreased towards R stage, as previously reported (Renato et al., 2014). Parameters V_t_ and τ_a_ were then used to determine the *in vivo* activities of COX and AOX pathways (Del-Saz et al., 2017). The *in vivo* COX pathway activity (ν_cyt_) remained unchanged from MG to BR stage and later decreased, being significantly lower in R tomato fruits (Fig.1B). On the other hand, the *in vivo* AOX pathway activity (ν_alt_) significantly (P<0.05) increased at the BR stage becoming the main contributor to climacteric respiration. These results underline the critical importance of assessing *in vivo* AOX and COX activities, as determined by ^18^O discrimination analysis.

### Transcript and metabolite changes in WT and *ghost* tomato fruits during ripening

Before assessing the metabolic profiles of WT and *ghost* fruit during ripening, we used qRT-PCR analysis to confirm that the color-based classification of WT and *ghost* fruits (Fig. 1A) corresponded with similar ripening stages. Ripening marker genes such as *E8* (encoding 2-oxoglutalate-dependent dioxygenase), *ACS2* (aminocyclopropane-1-carboxylate synthase 2), and *PG2a* (polygalacturonase 2a) displayed very low or undetectable expression at the MG (WT) or MW (*ghost*) stages, whereas a prominent increase was observed at the BR stage, when climacteric ethylene and respiration responses are induced (Fig. 1C). While *ACS2* transcript levels kept increasing at later stages of ripening, the expression of *PG2a* and, to a lower extent, *E8*, decreased in ripe fruit at the R (WT) or Y (*ghost*) stage (Fig. 1C). These patterns resemble those previously reported in tomato fruits (Guo et al., 2017)and validate the adequate sampling and selection of the different ripening stages in both WT and *ghost* fruits.

Metabolic profiles of WT and *ghost* lines during ripening were determined at the same fruit samples like those used for qRT-PCR analysis. A total of 48 and 47 primary metabolites were annotated after GC-TOF-MS analyses in fruit pericarp tissues from WT and *ghost* plants, respectively (Fig. 2, Supplemental Table S2, S4 and S5). Relative metabolite levels were determined after normalization by the mean levels of WT and *ghost* pericarp samples at MG and MW stages, respectively (Fig. 2, Supplemental Table S4 and S5). The levels of most detected metabolites significantly changed (P<0.05) at the onset of ripening (BR stage), typically showing similar levels or trends at later ripening stages (Fig. 2A). Considering the observed increase in AOX respiration at the BR stage (Fig. 1D), we focused our analysis on the significant changes observed at the BR stage (Fig. 2B, Supplemental Fig S3). The complete metabolite profiling data set and statistical analyses are shown in Supplemental Tables S4 and S5. Most of the metabolites displaying significantly lower levels at BR stage in WT fruit were amino acids including valine (Val; 0.22-fold), isoleucine (Ile, 0.50), glycine (Gly; 0.35), proline (Pro; 0.49), alanine (Ala; 0.41), threonine (Thr; 0.66), tyrosine (Tyr; 0.40) and tryptophan (Trp; 0.48), but also glycerate (0.65), beta-alanine (0.41) and raffinose (0.46) (Fig. 2B, Supplemental Table S4). The only two amino acids displaying significantly higher levels at the BR stage were aspartate (Asp; 1.79-fold) and methionine (Met; 1.75-fold). Other metabolites significantly (P<0.05) increased at the BR stage in WT fruits were dehydroascorbate (Dhasc; 2.29-fold), galacturonate (3.26), xylose (1.45), pyruvate (Pyr; 1.87), citrate (Cit; 1.74), 2-oxoglutarate (2-OG; 3.35), succinate (Succ; 2.37) and adenosine-5-monophosphate (2.29). Changes in primary metabolism detected in *ghost* tomato fruits showed a climacteric response in the absence of chromorespiration similar to that observed in WT fruits, although with some re-arrangements. Organic acids including Cit (2.36), 2-OG (2.49), Succ (2.70), Dhasc (1.34) and galacturonate (not detected at MW), as well as xylose (3.13) and adenosine-5-monophosphate (not detected at MW) increased in *ghost* fruits at the BR stage (Fig. 2A, Supplemental Fig. S3). Also as in WT fruits, the amino acids Met (3.83) and Asp (2.70) were increased, while Val (0.64-fold) and Ala (0.36) were decreased. However, other Asp family amino acids were only increased in *ghost* fruits, including Thr (2.70), homoserine (2.18) and asparagine (Asn; 3.77) as well as glutamate (Glu; 8.74), lysine (Lys; 2.15), Trp (6.64), ornithine (Orn; 2.19) and serine (Ser; 3.34). In addition, malate (0.60), *myo*-inositol (0.52), and GABA (0.63) displayed substantially lower levels in *ghost* BR fruits (Fig. 2A, Supplemental Fig. S3).

**Fig. 2.**
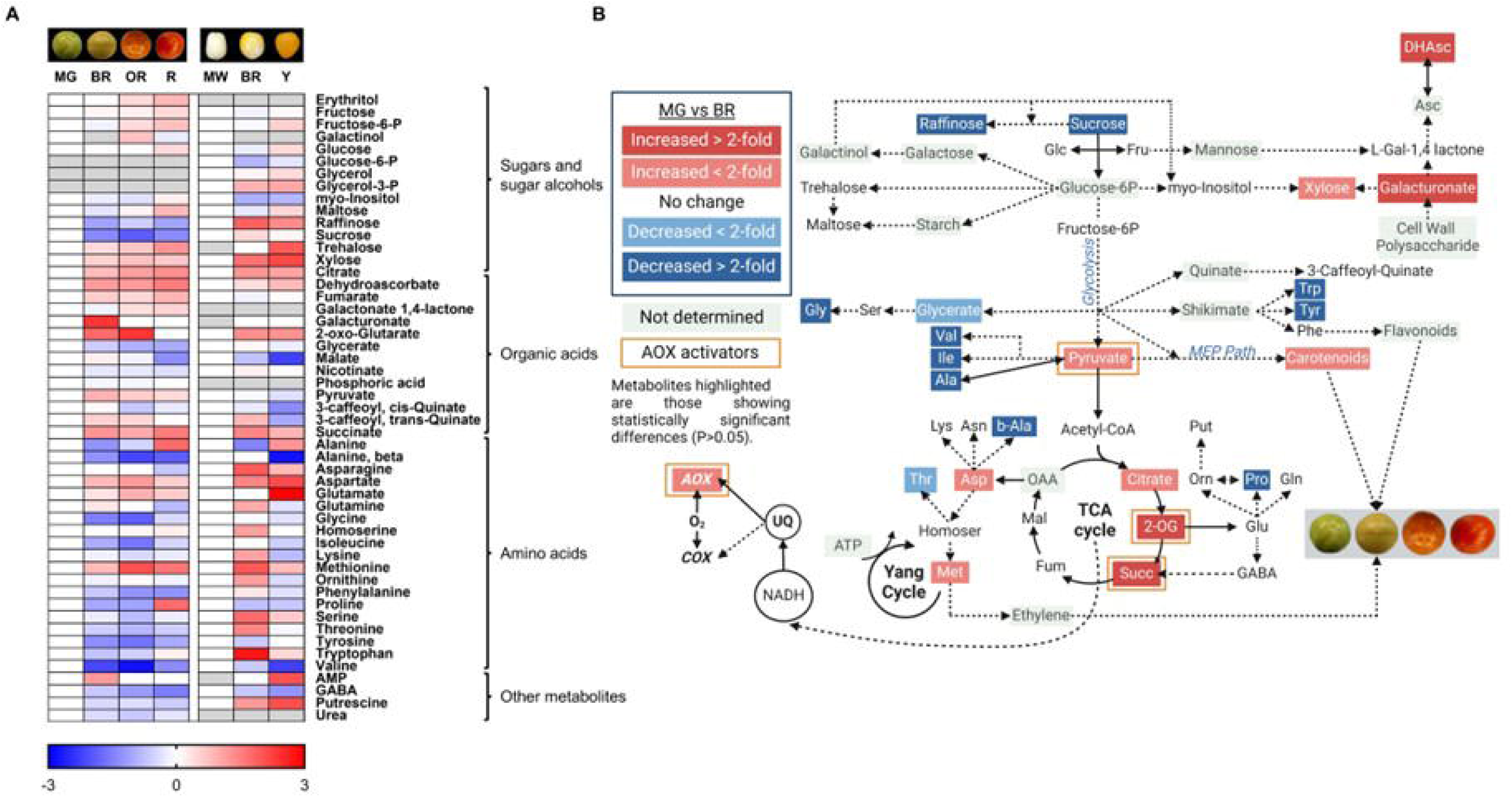
Metabolic profiles and model showing relative changes in primary metabolism in WT and *ghost* fruits at different ripening stages. (A) Heatmap showing the relative levels of metabolites analyzed by GC-MS in WT and *ghost* mutant fruits. Relative metabolite levels were normalized to the mean level of the MG (WT) or MW (*ghost*) stage and log2-transformed (i.e., levels at MG and MW stages are set to 0). Red and blue colors represent log2 fold-increases and decreases in metabolite levels, respectively. Gray indicates metabolites not detected under some experimental conditions. Values are means ± SE of 4–6 replicates. Statistical differences between MG or MW and other ripening stages are shown in Supplemental Tables S4 and S5. (B) Metabolic model showing changes in respiratory and associated metabolism at the climacteric peak of WT tomato fruit ripening (BR stage). Blue and red boxes indicate significantly (P < 0.05) decreased or increased metabolites and respiratory activities, respectively, in BR as compared to MG stage. Activators of the AOX1a (Pyruvate and 2-oxoglutarate) and AOX1b (succ) isoforms are highlighted with orange boxes. Image created with BioRender (https://biorender.com).

The metabolic profiling of WT and *ghost* fruit was complemented with the HPLC-DAD analyses of carotenoids (Supplemental Fig. S4). The biosynthesis of carotenoids, which is strongly activated in WT fruit from the BR stage, starts with the production of phytoene. Then, desaturation reactions that require PTOX activity in ripening fruit (Shahbazi et al., 2007) convert non-colored phytoene into red-colored lycopene. Further downstream reactions produce β-carotene (the main pro-vitamin A carotenoid) and lutein (the most abundant carotenoid in the chloroplasts of green tissues) (Rodriguez-Concepcion et al., 2018). An increase in the levels of phytoene, lycopene and β-carotene in concert with a decrease in lutein was observed during ripening of WT fruits (Supplemental Fig. S4). Also as was to be expected, lycopene became the most abundant carotenoid in R fruit (Supplemental Fig. S4). By contrast, phytoene was virtually the only carotenoid found in *ghost* fruit, reaching levels that were significantly (P<0.05) higher than those of lycopene and total carotenoids in WT fruits when ripe (Y vs. R).

### Generation and fruit development of AOX1a knock-out lines

The same RNA samples used for the analysis of ripening-related genes (Fig. 1C) were used to check the expression of AOX-encoding genes previously found to be up-regulated during tomato fruit ripening (Xu et al., 2012), *AOX1a* and *AOX1b*. Transcript levels of *AOX1a* steadily increased during fruit ripening and peaked at the R / Y stage (Fig. 1D). However, *AOX1b* expression did not significantly change even though they showed a tendency not to increase but to decrease during ripening (Fig. 1D). Based on these data, we decided to generate mutants with no AOX1a function to provide genetic evidence supporting the role of the AOX pathway during tomato fruit ripening. After transformation of MicroTom WT plants, two independent, Cas9 negative, homozygous lines were selected and named *a1.1* and *a2.3* (Fig. 3). The nucleotide deletions in both lines generated premature translation STOP codons (Fig. 3A), thus resulting in mutant proteins that were shorter (62 aa for *a1.1*, and 48 aa for *a2.3*) than the WT (358 aa). Therefore, we considered these two alleles as *aox1a* knockout mutants and selected them for the rest of the experiments. In agreement, AOX protein immunodetection (Fig. **3b**) and capacity measurements (Fig. 3C) in leaves from WT and *aox1a* mutant plants confirmed the effects of the *AOX1a* mutation. The AOX protein was only detected in WT samples (Fig. 3B) and both mutant lines displayed a strong reduction in AOX capacity compared to WT plants (Fig. 3C).

**Fig. 3.**
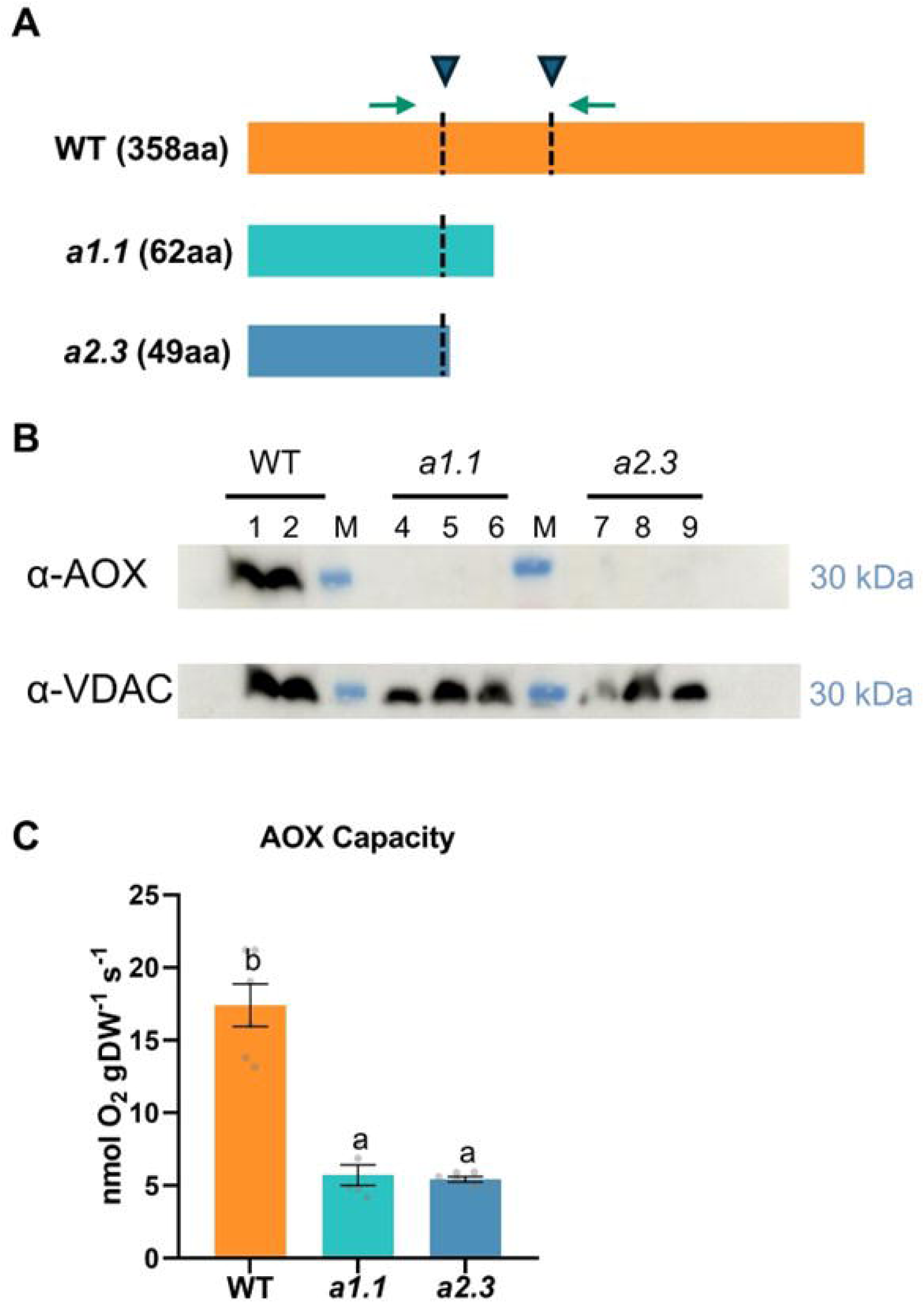
Tomato AOX1a-defective CRISPR-Cas9 mutants genotyping, AOX protein predictions, levels and capacity. (A) Scheme representing the predicted wild-type AOX1a protein, and the mutant versions generated by CRISPR-Cas9. The region targeted by the designed sgRNAs is indicated with an arrowhead and a dotted line. Green arrows represent the position of primers for PCR-based genotyping. (B) Western blot analysis of AOX and porin (voltage-dependent anion-selective channel protein 1–5, VDAC) protein levels from total leaf protein extracts (see section M&M section for details). (C) AOX capacity in leaves of WT and *aox1a* mutant lines measured as KCN-resistant respiration. Values are means ±SE of n=6 leaves from independent plants.

Regarding transition events from vegetative to reproductive stages, both *aox1a* mutant lines displayed a slight delay on the first fruit appearance as compared to WT plants (Fig. 4A, P=0.049 for *a1.1* and P=0.051 for *a2.3*). Despite this delay, the ability to reproduce and generate fruits was not impaired in the *aox1a* mutants. Nevertheless, the total number of fruits produced per plant was lower in both *aox1a* mutant lines (Fig. 4A), and their fruits displayed significantly (P<0.05) lower weight, height, and diameter (Fig. 4B).

**Fig. 4.**
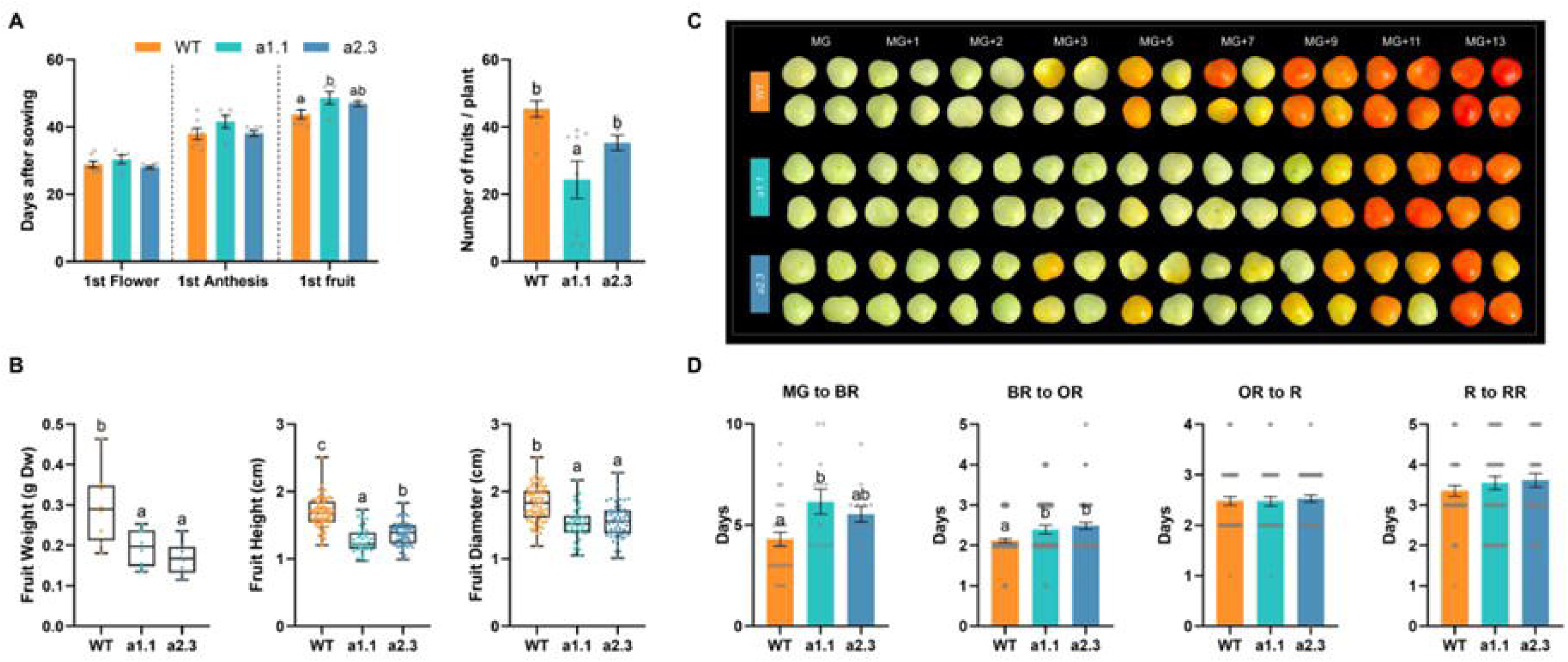
Tomato WT and *aox1a* mutant (*a1.1* and *a2.3*) fruits at different developmental stages. (A) Number of days to reach different developmental stages, and total number of fruits per plant. (B) Fruit diameter, dry weight, and height at red stage. Values are means ±SE of n=8 independent plants. Letters denote statistical differences (P<0.05). (C) Pictures of harvested tomato fruits at mature green (MG), MG+1, MG+2, MG+3, MG+5, MG+7, MG+9, MG+11, MG+13 stages. (D) Number of days from the MG to the breaker (BR) stage, BR to the orange stage (OR), OR to the red (R) stage, and R to the red ripe (RR) stage. Values are means ±SE of n=13-90 independent fruits. Letters denote statistical differences (P<0.05).

The *aox1a* mutant fruits also displayed a delay on the onset of ripening, as can be visually observed at MG+5, MG+7 and MG+9 stages when compared to WT fruits (Fig. 4C). Transition times from MG to BR and from BR to OR stages were significantly longer in both *aox1a* mutants compared to WT fruits (Fig. 4D). However, the subsequent transitions from OR to R and from R to Red Ripe (RR) stages were similar in all lines (Fig. 4D).

### Respiration and ripening-related gene expression in *aox1a* mutant fruits

The rates of total respiration and AOX capacity (V_alt_) were markedly altered in fruits of both *aox1a* mutant lines as compared to WT (Fig. 5A). Total respiration at MG stage was generally high in all three lines, probably driven by the energy demand for growth still present at our experimental conditions (Fig. 5A). Our intensive respiration measurements throughout several ripening stages revealed two respiratory peaks in WT fruits at the MG+2 (pre-BR) and MG+5 (BR) stages (Fig. 5A). After MG+5 stage, respiration rates steadily declined until R stages. Similar patterns of O_2_ consumption-based respiration rates were previously reported in tomato fruits (Xu et al., 2012; Renato et al., 2014), although only a single peak was reported. Both respiration peaks in fruits of the *a1.1* line were observed later (at MG+3 and MG+9 stages) compared to WT fruits, and only a single peak of respiration was evident (at the MG+5 stage) in fruits of the *a2.3* line. These shifts or absence of respiration peaks observed in fruits of both mutants (Fig. 5A) are in line with the observed delay at the onset of ripening (Fig. 4). In addition, V_alt_ was consistently and markedly lower in fruits of both *aox1a* mutants at all ripening stages as compared to WT fruits (Fig. 5A). Moreover, V_alt_ steadily increased from MG+3 to MG+5 in WT fruits while this response was absent in the two *aox1a* mutant lines (Fig. 5A). Although remaining under WT levels, V_alt_ was occasionally increased at MG+9 and MG+5 in *a1.1* and *a2.3* lines, respectively (Fig. 5A).

**Fig. 5.**
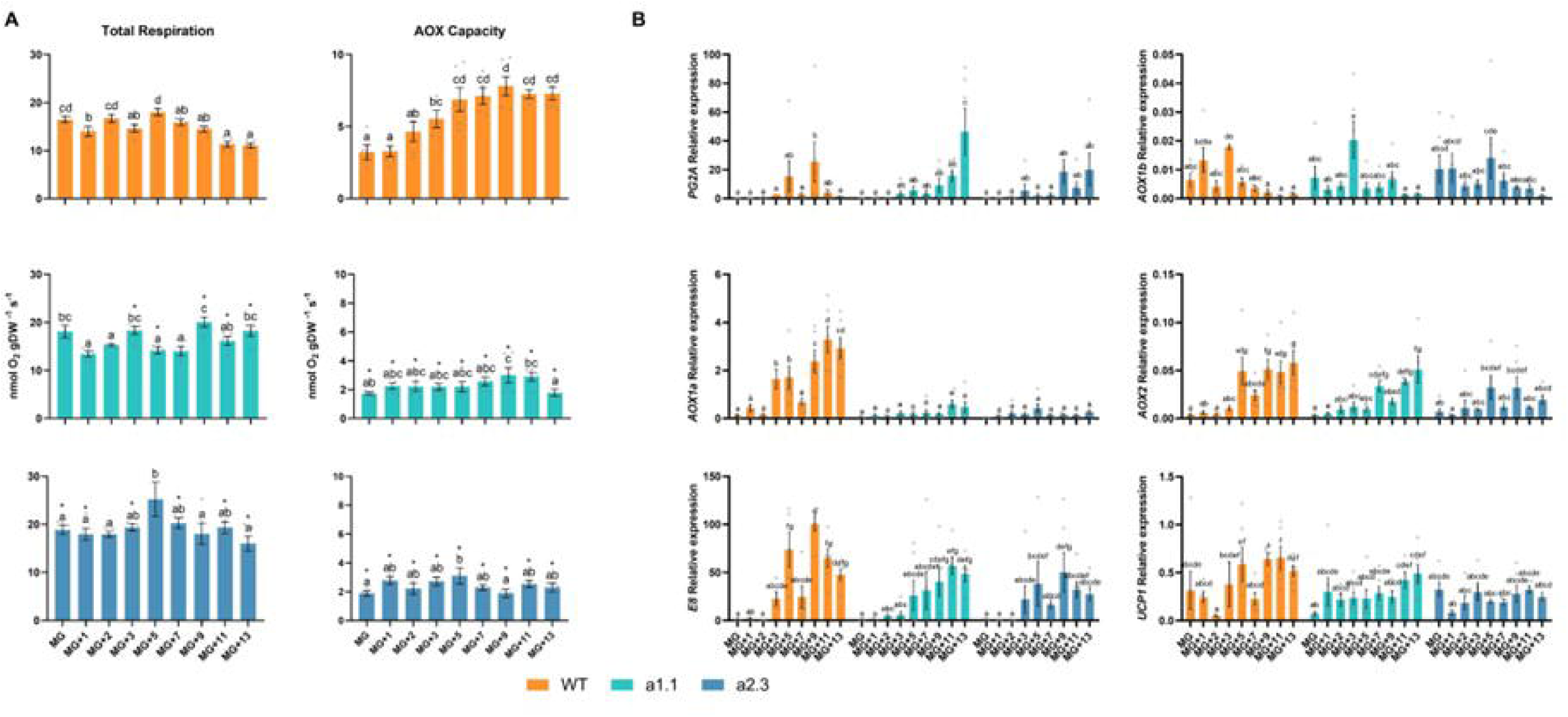
Respiration and gene expression analyses in fruits of WT and *aox1a* mutant lines at different ripening stages. (A) Total respiration and AOX capacity. Values are means ±SE of n=4-6 independent fruits. Letters denote statistically significant differences (P<0.05) between ripening stages in each line, and asterisks denote statistically significant differences (P<0.05) to WT samples at each ripening stage. (B) Relative transcript levels of AOX- and ripening-related genes in WT and *aox1a* mutant fruits at different ripening stages. AOX1a: alternative oxidase 1a; AOX1b: alternative oxidase 1b; AOX2: alternative oxidase 2; E8: 2-oxoglutarate-dependent dioxygenase ethylene responsive protein; PG2a: polygalacturonase 2a; UCP1: uncoupling protein 1. Values are means ± SE of 3-6 replicates. Significant differences (P < 0.05) are indicated by different letters.

In parallel to the observed changes in respiration, the expression of ripening-related genes was markedly altered in fruits of both *aox1a* mutant lines (Fig. 5B). The typical increase in transcript levels from ethylene-induced genes (*E8* and *PG2a*) was also delayed and/or attenuated in both *aox1a* mutants as compared to WT fruits (Fig. 5B). Therefore, our results strongly suggest that AOX1a is important for triggering the ethylene-induced responses in fruits during *on-vine* ripening. Regarding the expression of respiratory-related genes, different responses were observed in the *aox1a* mutants (Fig. 5B). The expression of the *AOX1a* gene in WT fruits followed a similar pattern to V_alt_ (Fig. 5B), thus suggesting a main role of the AOX1a isoform to determine the V_alt_ in tomato fruits. On the other hand, the expression pattern of *AOX2* and *UCP1* genes was similar in both *aox1a* mutants as compared to WT, although transcript levels were attenuated at some ripening stages, similarly as for the ethylene-induced E8 gene (Fig. 5B). However, this was not the case for AOX1b expression (Fig. 5B), which displayed disturbed patterns in *aox1a* mutants compared to WT.

### Primary and carotenoid metabolism in WT and *aox1a* mutant fruits

Profiles of primary metabolites were also characterized throughout ripening in fruits of WT and both *aox1a* mutant lines (*a1.1* and *a2.3*) (Fig. 6A). A clustered heatmap analysis clearly separated two groups of samples consisting of fruits at early and late ripe stages (Fig. 6A). This cluster separation supports previous results showing that the most profound changes in primary metabolite profiles during tomato fruit ripening occur from BR stage onwards (Osorio et al., 2011). In this line, the early pre-BR cluster included samples at MG, MG+1, MG+2, and MG+3 stages from all three lines, except for the case of MG+3 stage in the *a2.3* line (Fig. 6A). The subclades in this pre-BR cluster did not group the different lines, thus generally denoting metabolic consequences of the *AOX1a* mutation at early ripening stages were still modest. In contrast, the subclades of the cluster containing later ripening stages clearly separated WT from both *aox1a* mutant lines (i.e. from BR to end of ripening; MG+5, MG +7, MG +9, MG +11 and MG +13). The first subclade contained both mutant lines at the MG+13 stage, highlighting that the effects of *AOX1a* mutation on primary metabolism were accumulated consistently at the end of the ripening process. The second clade was further divided into three distinct subclades, two of them containing only mutant samples (except for WT at MG+7) and a third one separating WT samples (except for *a1.1* at MG+11). Primary metabolite data and statistical analyses are shown separately for early and late ripe clustered stages in Supplemental Tables S6 and S7, respectively. Among all these results, the primary metabolites displaying statistically significant changes between *aox1a* lines and WT are shown as bar plots separately at early and late ripe clustered stages in Supplemental Fig. S6 and S7, respectively. In general, this clustering pattern is in line with the delay on the transition around BR stage (Fig. 4) and denotes the important metabolic role of the AOX around BR stage, which finally generates a profound impact in primary metabolism at later stages.

**Fig. 6.**
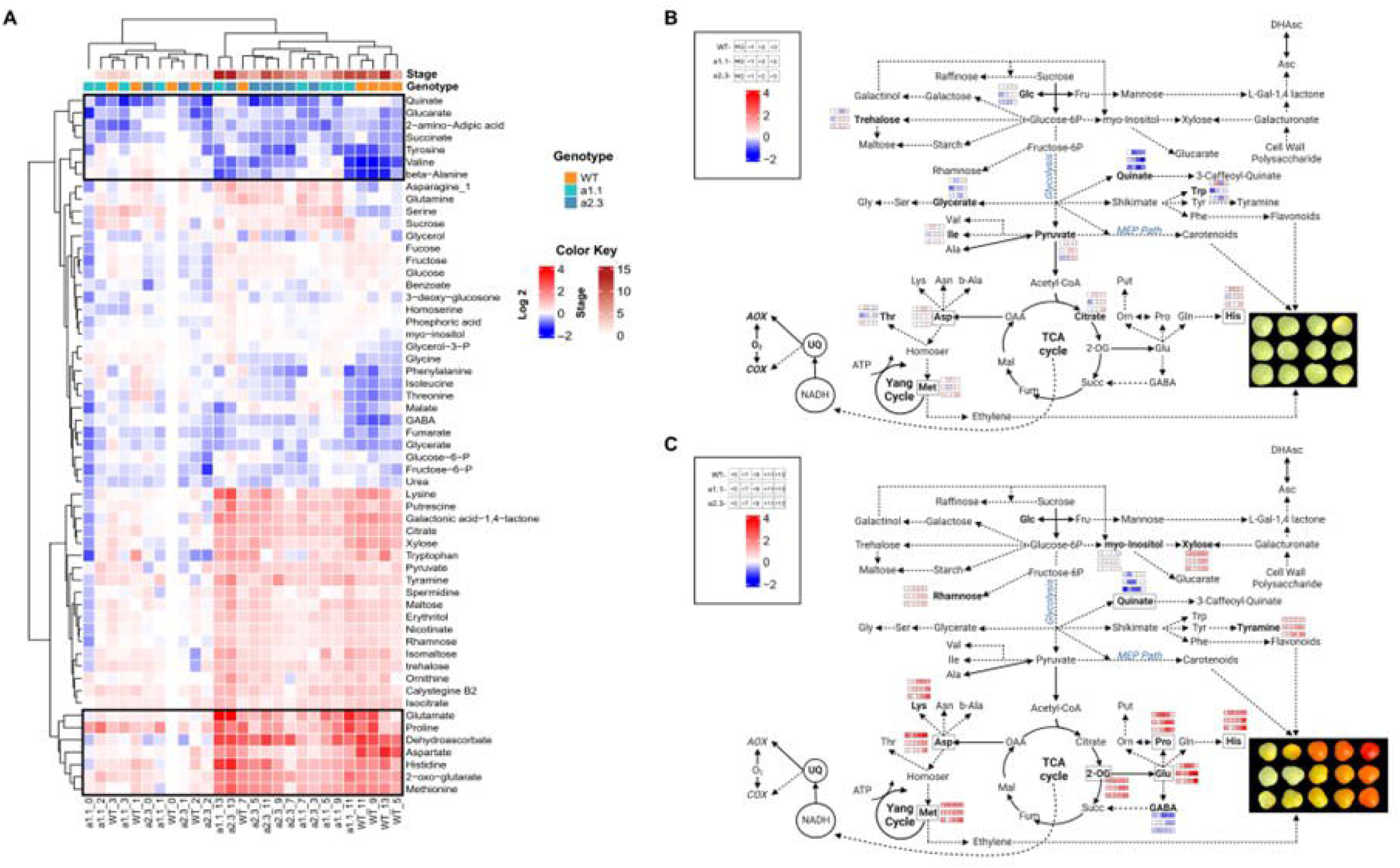
Metabolic profiles, cluster analysis and model showing relative changes in primary metabolism in fruits of WT and *aox1a* mutant lines at different ripening stages. (A) Clustered heatmap showing relative changes in primary metabolism in WT and *aox1a* mutant fruits at different ripening stages. Relative metabolite levels were normalized to the mean level of WT at the MG stage, and log2-transformed (i.e., levels at WT-MG are set to 0). Red and blue colors represent log2 fold-increases and decreases in metabolite levels, respectively. Cluster analysis was performed following by ‘ComplexHeatmap’ package in R. Dendrogram reveals correlation between tomato fruit lines, ripening stages and metabolites. Black boxes delimited the cases for which metabolites displayed the more pronounced responses (decreases at the top and increases at the bottom) during ripening. Values are means ± SE of 4–6 replicates. Statistical differences among lines at different ripening stages are shown in Supplemental Table S6 and S7. Metabolic models in (B) and (C) show changes in primary metabolism in WT and *aox1a* mutants at early and late ripening stages, respectively. Metabolites in bold and showing the heatmap results are only those displaying statistically significant changes (see analysis in Supplemental Table S6 and S7) between WT and *aox1a* lines at least at one ripening stage and consistently in both mutant lines. Grey boxes delimited the metabolites displaying statistically significant changes that were also included at the black boxes in (A). Image created with BioRender (https://biorender.com).

Regarding metabolite clusters, two major clusters separated the metabolites displaying increases along ripening (at the bottom of Fig. 6A) from those displaying decreases and/or minor changes (at the top of Fig. 6A). The two clades at the top of the cluster grouped seven metabolites with the more pronounced decreases along fruit ripening including 2-amino-adipic acid, *beta*-alanine, glucarate, quinate, Succ, Tyr, and Val (highlighted by a black box in Fig. 6A). Most of these metabolites were reported to decrease during fruit ripening in previous studies (Carrari and Fernie, 2006; Osorio et al., 2011; Quinet et al., 2019), and also in our results from WT Ailsa Craig fruits (Fig. 2, except Succ). However, we could not detect any consistent and statistically significant (P<0.05) difference when both mutants were compared to WT at any developmental stage, except for quinate (Supplemental Tables S6 and S7). On the other hand, the cluster at the bottom of the heatmap displayed the most pronounced increases at ripened stages including 2-OG, Asp, Dhasc, Glu, His, Met, and Pro (Fig. 6A), in concordance with our results in WT Ailsa Craig tomato fruits (Fig. 2, except for Glu and Pro). In this bottom cluster, statistically significant (P<0.05) differences in the patterns of all these metabolites were observed between WT and *aox1a* mutant fruits at late ripening stages with the exception of DHasc (Fig. 6C and Supplemental Fig. S7).

In line with disturbed primary metabolite responses, the reduction of chlorophyll levels occurred more rapidly in WT compared to the mutant lines, suggesting a difference in the degradation processes (Supplemental Fig. S5). Chlorophyll reduction is a reverse process to total carotenoid accumulation typically observed during fruit ripening (Trudel and Ozbun, 1970). In WT fruits, phytoene and lycopene were detected at MG+5 and MG+7 whereas they were minor or undetectable in *aox1a* mutant fruits (Supplemental Fig. S5). Thereafter, phytoene and particularly lycopene were boosted at MG+9 in WT fruits while they remained very low in both mutants. These results could denote a limitation of carbon skeletons for the biosynthesis of carotenoids in *aox1a* mutants. Nevertheless, phytoene and lycopene concentrations exhibited a significant increase in *aox1a* mutant lines at MG+11 and MG+13 stages, ultimately achieving WT carotenoid levels by MG+13 (Supplemental Fig. S5).

## DISCUSSION

### *In vivo* regulation of fruit respiration and consequences for energy metabolism during tomato fruit ripening

Respiration in fruits is essential to provide carbon skeletons and energy for the synthesis of nucleic acids and proteins as well as for carotenoids and derived flavor compounds, all being highly demanded during ripening (Tucker, 1993; Pott et al., 2019). In this context, the observed constant *in vivo* COP activity (ν_cyt_) up to the climacteric peak of respiration at the BR stage (Fig. 1B) could supply the ATP required for these high energy-demanding processes, particularly before the onset of ripening. Thereafter, the decrease of ν_cyt_ at R stage agrees with previous observations showing a decreased number of energized (ATP-producing) mitochondria in ripe fruits (Renato et al., 2014). On the other hand, the similar Δ_a_ values observed in WT and *ghos*t mutant fruits (Table 1) suggest a minor contribution of PTOX respiration to pericarp O_2_ consumption, although we cannot discard that ^18^O discriminations of PTOX (not directly determined here) and AOX could be similar. Together with previous evidence (Renato et al., 2014), our results suggest that chromorespiration could become of some relevance for ATP synthesis only at the end of ripening, when the *in vivo* COX respiration decreases. The lower mitochondrial ATP production in ripe fruits has been suggested to be influenced by increased expression of genes for AOX and UCP (Holtzapffel et al., 2002; Renato et al., 2014). However, our data suggest that decreased mitochondrial ATP production in ripe fruits is mainly a consequence of decreased ν_cyt_, rather than increased *in vivo* AOX activity (ν_alt_) (Fig. 1B). Indeed, ν_alt_ significantly (P<0.05) increased at the BR stage becoming the main contributor to climacteric respiration. These results agree with previous evidence suggesting that the AOX pathway is highly relevant during climacteric respiration in tomato (Xu et al., 2012; Colombié et al., 2017). Constraint-based metabolic modeling predicted an increase in fluxes through AOX and UCP respiration at the climacteric peak (Colombié et al., 2017). To this respect, our experimental data partly support these previous simulations and provide quantitative *in vivo* fluxes through both AOX and COX pathways during fruit ripening. The fact that ν_cyt_ was not increased during the climacteric peak could also reflect a minor role of the UCPs at this stage. Changes in UCP activity should impact the activity of the ATP-coupled COX pathway, as previously discussed in thermogenic tissues of the sacred lotus (Watling et al., 2006). Since direct measurements of the *in vivo* UCP activity are not currently plausible, further experiments involving UCP activity manipulations would be required to clarify the precise roles of the UCPs during fruit ripening.

Additional evidence on the participation of the AOX pathway during climacteric respiration is clearly provided here by our intensive respiratory analyses in newly generated KO *aox1a* mutants (Fig. 3 and 5A). Fruits of the *aox1a* mutants displayed shifts or absence of respiration climacteric peaks (Fig. 5A), which were in line with the observed delay of the onset of ripening and related gene expression (Fig. 4 and 5B). In agreement, delayed ripening and reduced total respiration have been reported in tomato fruits of AOX-silenced plants (Xu et al., 2012). Although ν_alt_ was not determined in previously reported AOX-silenced plants or in the currently obtained *aox1a* mutants, our results shown in Fig. 1B strongly suggest that any change on climacteric respiration observed was entirely due to changes on ν_alt_. Therefore, altered respiratory responses in our *aox1a* mutants are very likely due to changes in ν_alt_. Moreover, the consistent and strong reduction of AOX capacity (V_alt_) in fruits of both *aox1a* mutants at all ripening stages suggests that ν_alt_ response was limited in *aox1a* mutants at the time when WT fruits displayed their respiration peaks (MG+2 and MG+5; Fig. 5A).

Good correlations between AOX capacity and expression have been reported in several studies (reviewed in McDdonald et al., 2002 and in Del-Saz et al., 2018). However, total respiration patterns (i.e. increased until MG+5 and then decreased) were not correlated with V_alt_ or *AOX1a* expression, likely due to post-translational regulation mechanisms that are frequently reported to be the most important factors regulating respiration *in vivo* (Del-Saz et al., 2018). In line with this, the expression of *AOX1a* and *AOX1b* genes, which encode the main isoforms expressed in tomato fruits, did not correlate with ν_alt_ (Fig. 1B and 1C). AOX is usually found in its reduced state *in vivo* (Florez-Sarasa et al., 2019), which is a prerequisite for its allosteric activation by α-ketoacids through the formation of a thiohemiacetal bond occurring at conserved Cys residues. *In vitro* studies have shown that the activation of AOX by different organic acids from the TCA cycle is isoform-specific (Selinski et al., 2018). When the regulatory Cys position is substituted by a Ser residue by site-directed mutagenesis, the Ser-type AOX isoform becomes insensitive to 2-oxo acids (i.e. pyr and 2-OG) but is activated by succ (Selinski et al., 2018). This Ser-type AOXs occur naturally in some species (Grant et al., 2009) including the tomato isoform AOX1b (Holtzapffel et al., 2003). In the present study, an increase in key allosteric regulators of the Cys-type AOX1a (pyruvate and 2-oxoglutarate) and of the Ser-type AOX1b (succinate) were observed at the BR stage in concert with the increase of ν_alt_ (Fig. 1B and 2B). Together, these results strongly suggest a post-translational activation of the AOX at the BR stage through organic acids interaction. Notably, the fact that Ser-type AOX1b cannot be inactivated by Cys-oxidation suggests that this isoform can be highly active, if expressed, when succinate levels accumulate. While the level of AOX1b expression is lower than AOX1a, its special regulatory feature makes this isoform a possible candidate for (co-)regulating climacteric respiration together with AOX1a.

### Coordinated changes in primary metabolism and electron partitioning to AOX allows for the accumulation of key ripening-related metabolic precursors

In addition to the changes in TCA cycle intermediates, the coordinated increase of □_alt_ with other changes in key primary metabolites at the climacteric peak also suggests relevant metabolic roles of AOX (Fig. 2B). The combined decrease in sucrose and increase in pyr levels observed in WT fruits suggest a high glycolytic flux at the BR stage (Fig. 2B). Increased levels of pyr and other glycolytic intermediates can also serve as precursors for the synthesis of secondary metabolites such as flavonoids and carotenoids (Aharoni and Galili, 2011; Rodríguez-Concepción and Boronat, 2015; Zhang et al., 2015; Enfissi et al., 2017). In our study, altered patterns of glucose, glycerate and pyruvate accumulation were observed in *aox1a* mutants at early ripening stages (Fig. 6B, Supplemental Fig. S6 and Table S6), which might denote an alteration on the glycolytic pathway. While carbon flux experiments would be required to precisely determine quantitative changes in carbon partitioning, our data suggest that restriction on respiratory metabolism could lead to a limitation on the provision of carbon skeletons for the carotenoid pathway. Specifically, pyruvate together with glyceraldehyde 3-phosphate feed the methylerythritol 3-phosphate (MEP) pathway, which produces the prenyldiphosphate precursors required to synthesize for geranylgeranyl diphosphate (GGPP) and downstream carotenoids (Supplemental Fig. S4) (Rodriguez-Concepcion et al., 2018). Increased levels of pyruvate were consistently observed in ripe fruits of GGPP synthase mutants, which displayed an important restriction in carotenoid synthesis (Barja et al., 2021). In line with this, low levels of the carotenoids phytoene and lycopene were detected in *aox1a* mutants at stages when WT fruits started to boost their production (Supplemental Fig. S5). As commented above, *aox1a* mutants also displayed increased respiration rates, albeit displaced to later developmental stages, probably due to an increased contribution of the AOX1b isoform. This compensation may have allowed the *aox1a* mutants to achieve WT levels of carotenoids at the end of the ripening (Supplemental Fig. **S5**) and reach a similar color (Fig. 4C) as WT fruits by the MG+13 stage.

Importantly, the only two amino acids displaying high levels at the climacteric peak of AOX respiration were aspartate and methionine, which are precursors for the synthesis of ethylene (Fig. 2B), the most important hormone triggering fruit ripening. Respiration is thought to provide the ATP required for the Yang cycle, thus allowing the recycling of methionine for the biosynthesis of ethylene (Fig. 2; Yang and Hoffman, 1984). According to this, an increase in COX-dependent respiration would be expected at the climacteric peak given its ATP-coupled nature. However, our results clearly show that climacteric peak is mainly driven by the non-phosphorylating AOX pathway (Fig. 1B). Previous studies in leaf and root tissues suggest different roles of the AOX in providing metabolic flexibility by allowing the provision of demanded carbon skeletons under high cell energy (ATP) charge (Del-Saz et al., 2018). In this line, a high AOX respiratory flux at the climacteric peak would enhance the provision of key carbon precursors required for triggering fruit ripening. Despite the Yang cycle allows for the synthesis of ethylene precursors while recycling Met, there is evidence indicating that ‘*de novo*’ synthesis of methionine becomes relevant when high rates of ethylene production occur in tomato fruits (Katz et al., 2006). Noticeably, about 50-80% of the methionine produced in plants is used for the synthesis of ethylene (Ranocha et al., 2001; Hesse et al., 2004). Therefore, a high carbon flux towards methionine synthesis should be expected during the climacteric peak of ethylene production and could presumably be fulfilled from TCA cycle intermediates and aspartate (Fig. 2B). In close agreement, the accumulation of Asp in both *aox1a* mutant lines was almost abolished, while this metabolite displayed >10-fold increase at MG+5 in WT fruits, i.e. around BR stage, and reached even higher levels at the end of ripening (Fig. 6C and Supplemental Fig. S7). Given that respiration and V_alt_ was restricted in the *aox1a* mutants around BR stage (Fig. 5), limited mitochondrial NADH reoxidation probably hindered the activity of the TCA cycle dehydrogenases required for the provision of oxalacetate (precursor of Asp). Indeed, restricted activity of 2-oxoglutarate dehydrogenase could be the cause of the observed 2-oxoglutarate accumulation in the *aox1a* mutants, which is a key carbon precursor that links TCA cycle with amino acid metabolism through glutamate metabolism (Forde and Lea, 2007). Under high NADH matrix concentrations, the conversion of 2-oxoglutarate to glutamate would be favored in the *aox1a* mutants either via glutamate: 2-oxoglutarate aminotransferase (GOGAT) or glutamate dehydrogenase (GDH) (Forde and Lea, 2007), and against the generation of downstream TCA cycle intermediates such as oxalacetate. In agreement, Glu was very highly accumulated at the end of ripening in both mutant lines, while this behavior was absent in WT fruits (>50-fold change, Fig. 6C and Supplemental Fig. S7). Proline can be synthesized from glutamate (Kavi Kishor et al., 2022) and both metabolites displayed mirrored changes in WT and in both mutants, (Fig. 6C and Supplemental Fig. S7). Our results then denote that restricted AOX respiration in *aox1a* mutants greatly affected aspartate accumulation, thus limiting methionine synthesis which in turn could curtail ethylene synthesis. Although not directly determined here, ethylene production rapidly triggers an increase in the expression of ripening-related genes (i.e. *E8* and *PG2a*), which were delayed and/or attenuated in *aox1a* mutants as compared to WT fruits (Fig. 5B). Consistently, ripening of *aox1a* mutant fruit was slower that WT controls (Fig. 4). Moreover, our intensive sampling and analyses showed first increases of respiration (MG+2, Fig. 5A), V_alt_ (MG+3, Fig. 5A), *AOX1a* (MG+3, Fig. 5B) and *AOX1B* (MG+3, Fig. 5B) expression in WT fruits during pre-BR stages, which preceded the notable increases in ethylene responsive genes (MG+3 and MG+5, Fig. 5B). All these responses were clearly attenuated and/or shifted in the *aox1a* mutants (Fig. 5A and 5B), in parallel to the absent increase in threonine, methionine and aspartate during the pre-BR stage (Fig. 6B and Supplemental Fig. S6). Overall, respiration, metabolic profiling and expression analysis in the *aox1a* mutans and WT lines reveal the key role of the AOX pathway for the synthesis of aspartate family amino acids including aspartate and methionine, which are required for ethylene synthesis during climacteric ripening.

## Concluding remarks

We propose a positive feedback regulation by which increased pyruvate, 2-oxoglutarate and succinate levels produced from sucrose oxidation at the BR stage can allosterically activate AOX1a (and AOX1b), which allows the reoxidation of NAD(P)H escaping the ATP respiratory control. Such a climacteric burst of the non-phosphorylating AOX pathway respiration allows for a high rate of glycolytic and TCA cycle activities to provide key ripening-related metabolic precursors, such as aspartate and methionine for ethylene production and pyruvate for carotenoid synthesis. The restrictions in respiratory metabolism in AOX1a-defective CRISPR-Cas9 mutants strongly suggest the crucial role of the AOX pathway in triggering climacteric ripening by supplying of aspartate and methionine for the biosynthesis of ethylene. New opportunities are now open for measuring *in vivo* respiratory activities in climacteric and non-climacteric fruits as well as in ripening mutants, which will be key to better understanding fruit ripening and disentangle the precise hierarchy of the metabolic processes involved.

## MATERIAL AND METHODS

### Plant material and growth conditions

Seeds from tomato (*Solanum lycopersicum*) wild-type (WT) plants from the ‘Ailsa Craig’ and ‘MicroTom’ varieties as well as two CRISPR Cas9 mutant lines (*a1.1* and *a2.3*) were sown in small pots and grown under controlled walk-in-growth chamber conditions: 16h/8h light/dark photoperiod, day/night temperature 24°C /22°C; photosynthetic photon flux density (PPFD) of 350-400 μmol m^-2^ s^-1^. After two weeks, plants were transplanted into 2 L (cv. Ailsa Craig) or 0.5 L (cv. MicroTom) pots containing a mixture of substrate:vermiculite:perlite (3:1:1). Plant growth, fruit sample collection and fruit development analyses were carried out as described (Supplemental Methods S1). Each harvested fruit was cut into pericarp slices and divided into subsamples that were used for respiration analysis (fresh tissue) or immediately frozen in liquid nitrogen and stored at −80°C for subsequent RNA and metabolite analysis.

For the generation of the *aox1a* mutant plants, two single guide RNAs (sgRNA) sequences were designed to create short deletions using the CRISPRP 2.0 online tool (http://crispr.hzau.edu.cn/CRISPR2/; (Liu et al., 2017) shown in Fig. 3A. Cloning, transformation and *in vitro* regeneration of mutant plants was performed as described (Supplemental Methods S2). Two (*a1.1*) and (*a2.3*) stable T3 lines were used for further experiments, which were predicted to generate two different truncated AOX1a proteins (Fig. 3).

### Respiration and oxygen isotope discrimination analysis

Respiration and ^18^O discrimination analysis in pericarp fruit tissues required a set-up that is detailed in Method S3. Measurements of total respiration and AOX capacity in pericarp fruit tissues were performed by using liquid-phase Clark-type oxygen electrodes (Rank Brothers LTD Dual Digital Model 20) as described (Method S3). Thereafter, ^18^O discrimination analyses during respiration were performed by using a dual-inlet isotope ratio mass spectrometer (DI-IRMS) system as described (Supplemental Methods S3). After both electrode and DI-IRMS measurements, samples were oven-dried at 60 °C for at least 48 h to determine dry weights (DWs). All respiration and ^18^O discrimination data are means ± SE of 3-6 biological replicates corresponding to different fruits.

### Gene Expression analysis

RNA isolation, cDNA synthesis, and RT-qPCR analyses were carried out as described (Supplemental Methods S4). Tomato ACT4 (Solyc04g011500) was used as a reference gene to correct for differences in the total amount of transcripts and the 2-ΔΔCt method (Livak and Schmittgen, 2001) was used to calculate the fold-change of gene expression. Data were normalized to the mean value of MG and MW fruits for WT-Ailsa Craig and *ghost* mutants, respectively (i.e. the level of all transcripts for WT fruits at MG and *ghost* mutant fruits at MW were set to 1). For experiments in MicroTom WT and *aox1a* mutants, data were normalized to the mean value of WT tomato fruit at MG stage (i.e. the level of all transcripts for MicroTom WT fruits at MG was set to 1). Values presented in Fig. 1C and 5B are means ± SE of 3-6 biological replicates corresponding to different fruits.

### Western Blot Analysis

Protein extraction and western blot analysis was performed as previously described (Del-Saz et al., 2022). After protein extraction, SDS-PAGE gel electrophoresis and transfer to nitrocellulose membrane, diluted 1:500 polyclonal anti-AOX AOX1 and 2 (AS04054, Agrisera, Sweden) or diluted 1:5000 anti-Porin, voltage-dependent anion-selective channel protein 1–5 (AS07212, Agrisera, Sweden), were used as primary antibodies. After incubation with 1:20000 horseradish peroxidase secondary antibody (Cytiva) during 1h at room temperature, the detection of immunoreactive bands was performed using ECL SuperSignal™ West Femto Maximum Sensitivity Substrate (Thermo Fisher). Chemiluminescent signals were collected by IQ 800 Armersham (Cytiva). Two different immunoblot experiments per protein were performed with very similar results and the image shown in Fig. 3B belongs to one of the membranes obtained. Two and three samples corresponding to leaves from different WT and *aox1a* mutant plants at T3 generation, respectively, were used for the western Blot experiments (Fig. 3B).

### Primary metabolite and carotenoid profiling

Primary metabolite extractions were performed as described previously (Lisec et al., 2006) using approximately 10 mg of lyophilized pericarp tissue, previously frozen. Derivatization of all samples was carried out as described previously (Lisec et al., 2006), as well as the GC-TOF-MS analyses in WT-Ailsa Craig/*ghost* fruits. The GC-MS analyses in MicroTom WT and *aox1a* mutant fruits were carried out with a 5977C GC/MSD (Agilent) following parameters described previously (Lisec et al., 2006) with modifications on the MS scan rate. Metabolites were identified manually by TagFinder software (Luedemann et al., 2012) using the reference library mass spectra and retention indices housed in the Golm Metabolome Database (http://gmd.mpimpgolm.mpg.de) (Kopka et al., 2005). The parameters used for the peak annotation of all detected metabolites can be found in Supplemental Tables S2 and S3, which follows previously reported recommendations (Fernie et al., 2011). Data in WT-Ailsa Craig/*ghost* fruits were normalized to the mean value of MG and MW fruits for WT and *ghost* mutants, respectively (i.e., the value of all metabolites for MG or MW was set to 1). Data MicroTom WT and *aox1a* mutant fruits were normalized to the mean value of WT-MG fruits (i.e., the value of all metabolites for MicroTom WT fruits was set to 1). Values presented in Supplemental Tables S4 to S7 are means ± SE of 4-6 biological replicates corresponding to different fruits.

Carotenoids and chlorophylls were determined as previously described (Barja et al., 2021) by using the Agilent 1200 series HPLC system (Agilent Technologies).

### Statistical and data analyses

For the statistical analyses in Table 1, Fig. 1, 4, 5, Supplemental Fig. S1, S3, S5, S6, S7, Supplemental Table S6 and S7, a one-way ANOVA with a level of significance of P < 0.05 was performed with the SPSS statistical software package, version 25 (IBM Corp., 2016, Armonk, New York, NY, USA), and Duncan’s posthoc test was used to determine statistically significant differences. Student’s t-tests were used for the statistical analyses in Fig. 2B, Supplemental Fig. S2, S4, Supplemental Tables S4 and S5 to determine significant (P < 0.05) differences between MG and other ripening stages. Finally, Student’s t-test with a level of significance of P < 0.05 were also applied in Fig. 5A to compare the *aox1a* mutant lines against WT plants at each ripening stage.

## ACKNOWLEDGEMENTS

We would like to thank Biel Martorell and Maria Rosa Rodriguez-Goberna for their technical help while running DI-IRMS, GC-MS and HPLC analyses.

## AUTHOR CONTRIBUTIONS

IFS and Manuel RC designed the research; AIS, NFDS, ME, EFP and IFS performed the research; Miquel RC and ARF contributed analytic tools; AIS, NFDS, ME, EFP, IFS and Manuel RC analyzed data; AIS, IFS and Manuel RC wrote the manuscript. All authors revised and approved the manuscript.

