## Supplementary Material for "Activation of alternative oxidase ensures carbon supply for ethylene and carotenoid biosynthesis during tomato fruit ripening"

**SUPPLEMENTAL INFORMATION**

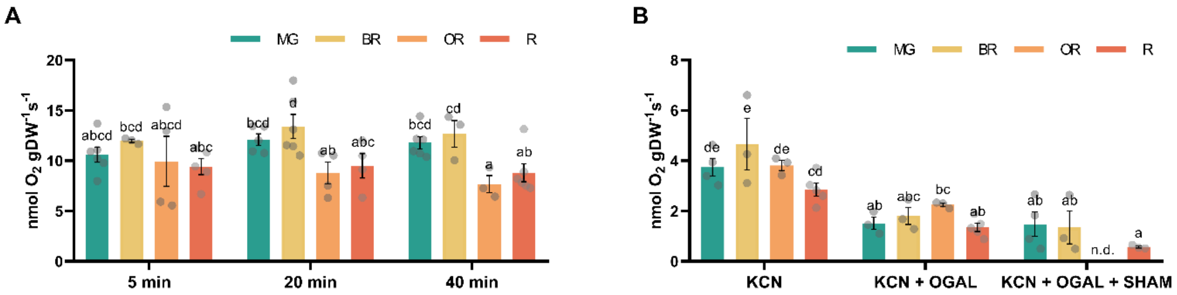


**Fig. S1 Respiratory pathways in wild-type (WT) fruits at different ripening stages** (A) Total respiration rates after 5, 20, and 40 minutes of incubation with respiration buffer. (B) Respiration rates after sequential additions of 5 mM potassium cyanide (KCN), 1 mM octyl gallate (OGAL) and 20 mM salicylhydroxamic acid (SHAM). Values are means ± SE of 3 to 6 replicates. Significant differences ($P < 0.05$) are indicated by different letters. "n.d." denotes "not detected

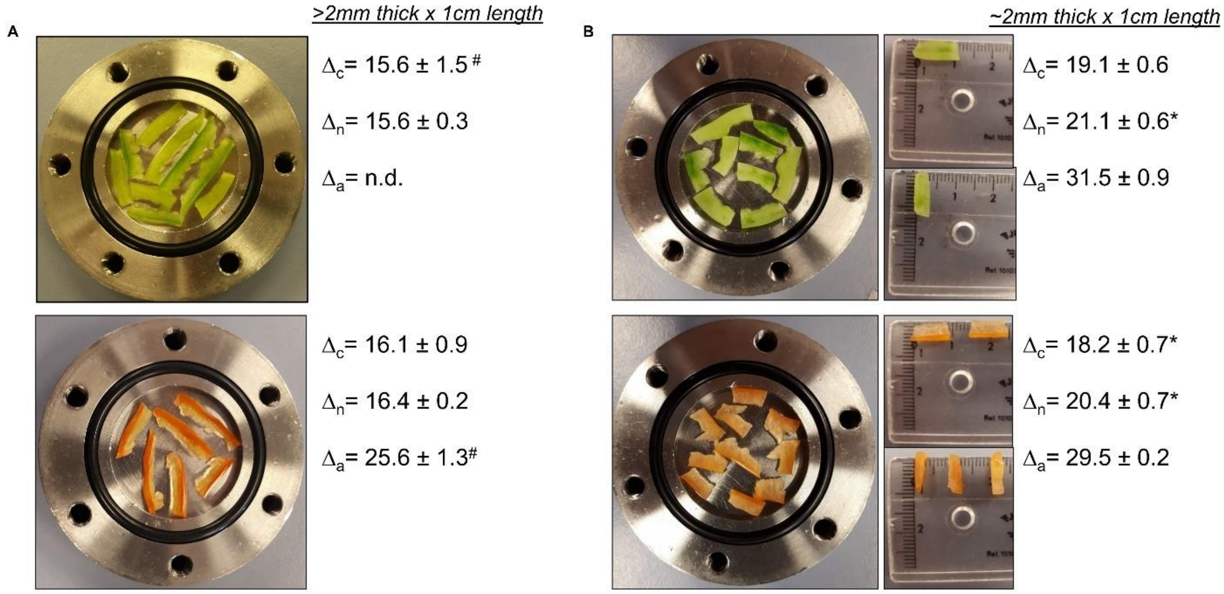


**Fig. S2 Photographs of the sliced pericarp tissue at MG and R stages of different sizes placed at the DI-IRMS cuvette, and effects on the 18O discrimination.** Pericarp slices of more than 2 mm thick and 1 cm length (A), and of approx. 2 mm thick and 1 cm length (B) are shown together with their corresponding values of 18O discrimination by the AOX pathway (Δa, after 10mM KCN treatment), by COX (Δc, after 20mM SHAM treatment), and in the absence of inhibitors (Δn). The 18O discrimination data are means ± SE of 3-5 biological replicates corresponding to different fruits. Asterisks denote significant differences ($P < 0.05$). n.d., 'not determined'; #only 2 replicates available**.**

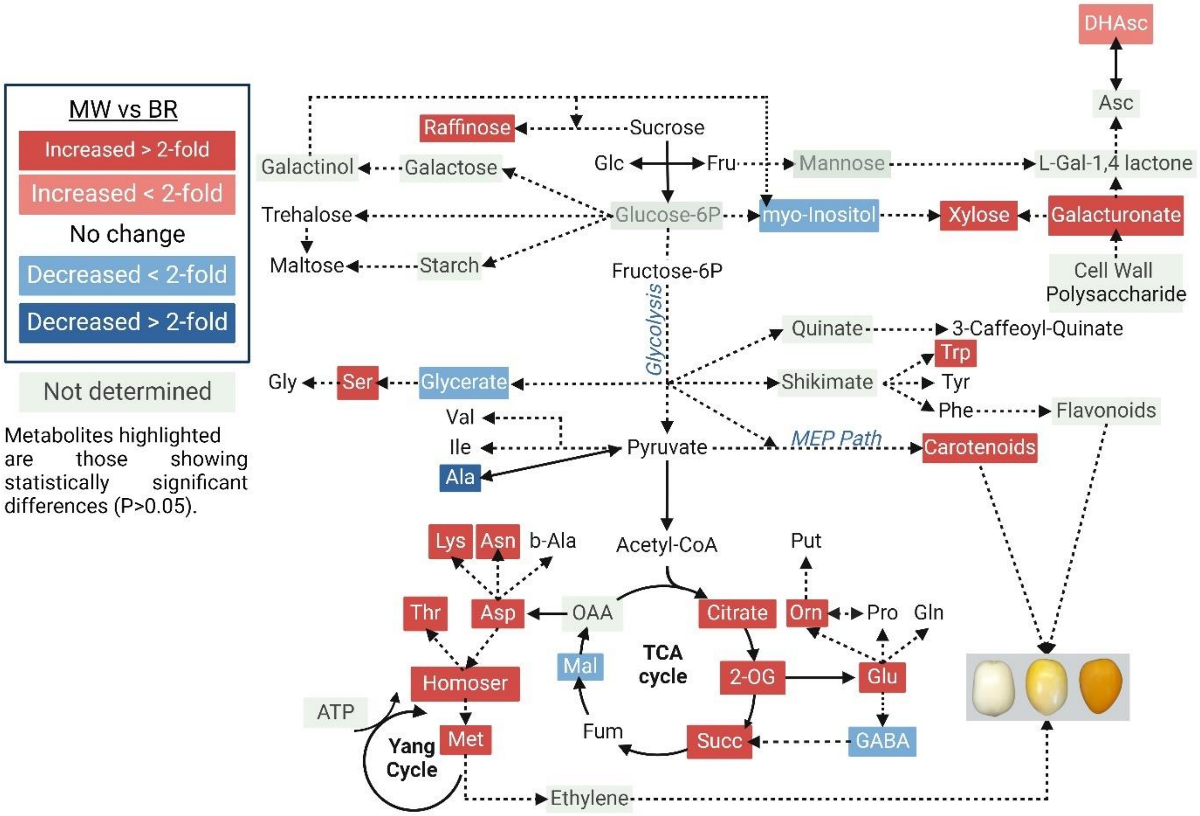


**Fig. S3 Changes in respiratory and associated metabolism at the climacteric peak of *ghost* tomato fruit ripening (BR stage).** Blue and red boxes indicate significantly (P < 0.05) decreased or increased metabolites and respiratory activities, respectively, in BR as compared to MW stage. Image created with BioRender. (https://biorender.com).

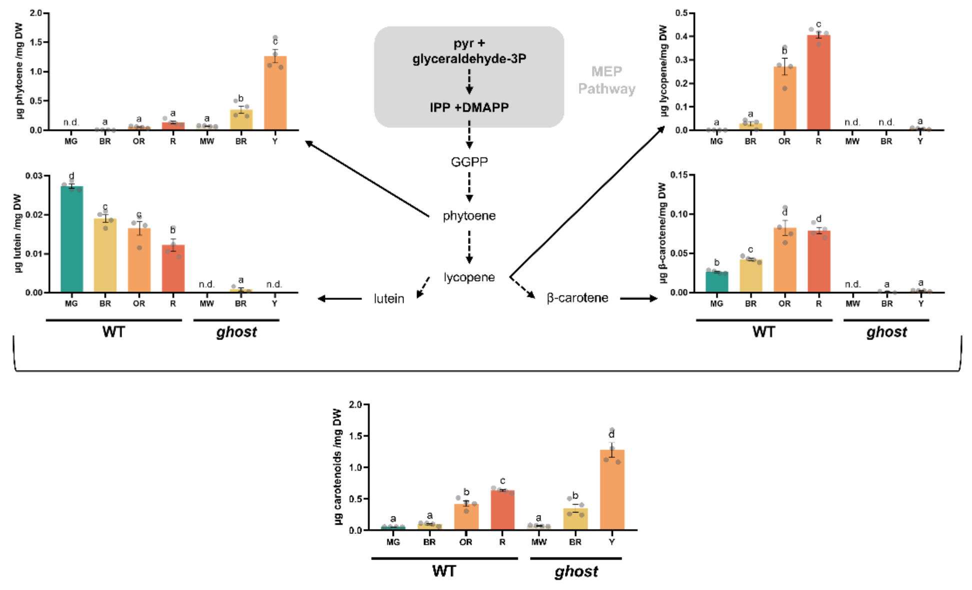


**Fig. S4 Levels of individual carotenoids (phytoene, lycopene, lutein and b-carotene) and total carotenoids in fruits of WT and *ghost* mutant at different ripening stages.** Values are means ± SE of 4 replicates and different letters represent statistically significant differences (one-way ANOVA followed by Duncan's multiple comparisons test, $P < 0.05$).

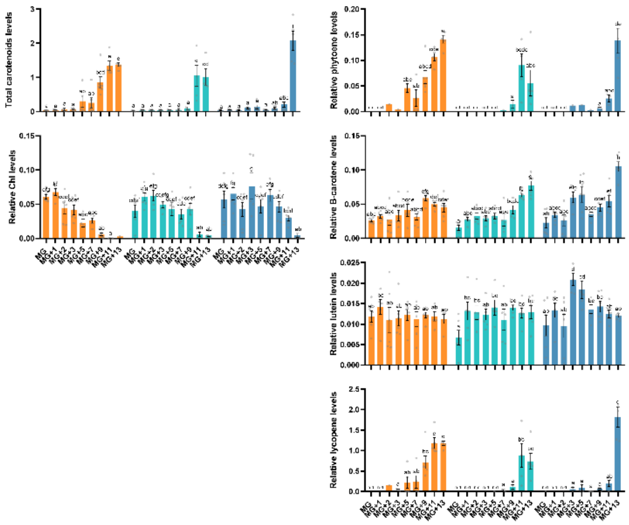


**Fig. S5 Levels of individual carotenoids (phytoene, lutein, lycopene, and b-carotene) as well as total carotenoids and chlorophylls in fruits of WT and *aox1a* mutant lines at different ripening stages.** Values are means ± SE of four to six replicates and different letters represent statistically significant differences (P < 0.05).

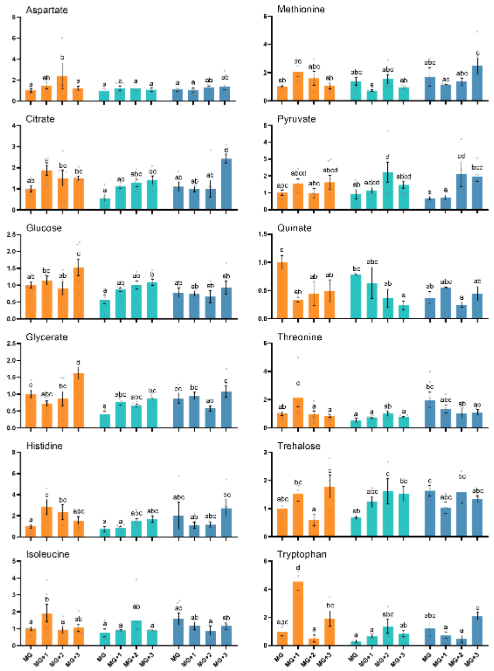


**Fig. S6 Relative levels of individual metabolites that displayed statistically significant (P<0.05) differences between WT and *aox1a* mutant fruits during early ripening stages.** Values are means ± SE of four to six replicates and different letters represent statistically significant differences (P < 0.05).

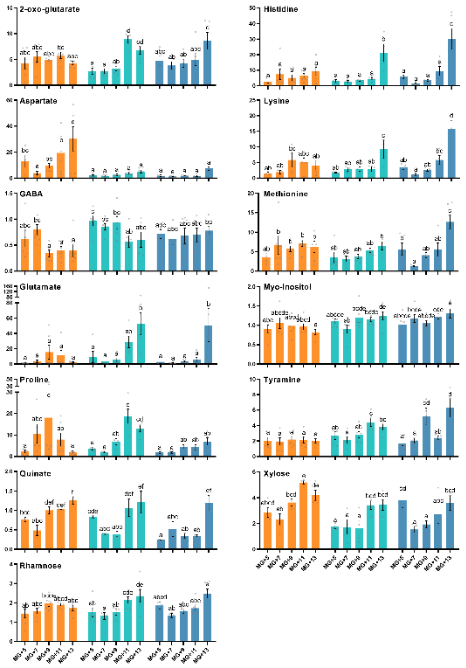


**Fig. S7 Relative levels of individual metabolites that displayed statistically significant (P<0.05) differences between WT and *aox1a* mutant fruits during late ripening stages.** Values are means ± SE of four to six replicates and different letters represent statistically significant differences (P < 0.05).

**Table S1  Primers used in this work**

| Name | Gene | Primer Name | Sequence 5'-3' |
|---|---|---|---|
| Cas9 | | Cas 9-Fw | TCCCTCATCAGATCCACCTC |
| | | Cas 9-Rv | CTGAAACCTGAGCCTTCTGG |
| RNA-guide 1 | Solyc08g075540 | RNA guide 1-Fw | ATTGCGTTACTTTTCAACTACAGT |
| | | RNA guide 1-Rv | AAACACTGTAGTTGAAAAGTAACG |
| RNA-guide 2 | Solyc08g075540 | RNA guide 2-Fw | ATTGGGAGAAAGTGGCAGTAACGT |
| | | RNA guide 2-Rv | AAACACGTTACTGCCACTTTCTCC |
| Sl-AOX1a | Solyc08g075540 | SlAOX1a-Geno-Fv | CCTTCTTCCTCAAGTTTCTTTC |
| | | SlAOX1a-Geno-Rv | CAGTACCATCTGGTTTAGTAG |
| Sl-AOX1a | Solyc08g075540 | SlAOX1a-qPCR-Fv | TTGTTTTAGGCCATGGGAGACTT |
| | | SlAOX1a-qPCR-Rv | CAGTGGGGAAACGAAGGATCT |
| Sl-AOX1b | Solyc08g075550 | SlAOX1b-qPCR-Fw | CACACTGATGCAACCAACGC |
| | | SlAOX1b-qPCR-Rv | TACGTCTCCCATGGCCTGAA |
| Sl-AOX1c | Solyc08g005550 | SlAOX1c-qPCR-Fw | CCACTTTGCATCGGATATTTATTAC |
| | | SlAOX1c-qPCR-Rv | AGTGATACCCAAGTGGAGCTG |
| Sl-AOX2 | Solyc01g105220 | SlAOX2-qPCR-Fw | CGCAAGTTCGAGCACAGT |
| | | SlAOX2-qPCR-Rv | TAGGCAGTCTCCAGTAGTCAAT |
| Sl-ACS2 | Solyc01g095080 | SlACS2-Fw | CGTTTGAATGTCAAGAGCCAGG |
| | | SlACS2-Rv | TCGCGAGCGCAATATCAAC |
| Sl-PG2a | Solyc10g080210 | SlPGA-qPCR-Fw | ATGGCAATGGACAAGTATGGTG |
| | | SlPGA-qPCR-Rv | TTAAGGCCGTTGGTGCATC |
| Sl-E8 | Solyc09g089580 | SlE8-qPCR-Fw | AGCTGCAAGTTGGAGAGACACG |
| | | SlE8-qPCR-Rv | CCGCATGGAGTTGGAAATTC |
| Sl-ACT4 | Solyc04g011500 | Sl ACT4-qPCR-Fw | CCTTCCACATGCCATTCTCC |
| | | Sl ACT4-qPCR-Rv | CCACGCTCGGTCAGGATCT |
| Sl-UCP1 | Solyc09g011920 | Sl UCP1-qPCR-Fw | TGATCAAGTGAAGGAGGCTGT |
| | | Sl UCP1-qPCR-Rv | AGCCCCGCAATCAAATGA |

**Table S2 Parameters used for peak annotation of all detected metabolites in WT-Ailsa Craig and *ghost* fruits.**

**Supplemental Table 2.** Overview of the metabolite reporting list.

**GC-TOF-MS metabolites**

| | A | B | C | D | E | F | G | H | I | H | I | H |
|---|---|---|---|---|---|---|---|---|---|---|---|---|
| 1 | **Experiment title:** | | | Metabolite profiles of tomato wild-type and ghost mutants during ripening | | Peak/compound no.- number referenced back to the main text | | | Mass to charge ratio (m/z) | | | |
| 2 | **Organism/Plant species:** | | | Solanum lycopersicum | | Ret . Time- Time expected, Tag Time Index and Time deviation | | | (S)- identification confirmed by a standard compound | | | |
| 3 | **Organ/tissue:** | | | Fruit Pericarp | | Putative Name- putative identification of the metabolite/derivative | | | I, II, III- different isomers | | | |
| 4 | **Analytical tool:** | | | GC-TOF-MS | | Corresponding metabolite name in literature | | | Identification level (A; B; C; D)- (A) standard or NMR; (B) MS/MS; (C) $MS^E$; (D) MS only | | | |
| 5 | | | | | | Mol. Formula- molecular formula of the metabolite or its FA adduct; | | | | | | |
| 6 | **Peak/ Compound no.** | **Time Expected** | **Tag Time Index** | **Time Deviation** | **Putative metabolite name (Derivative)** | **Corresponding Metabolite in Literature** | **Metabolite Class** | **Mol formula** | **Mass to charge ratio (m/z)** | **Species detected before** | **References** | **Identification level (A-D)** |
| 7 | 1 | 222650 | 222348 | -0,15 | M000071_A104002-101_METB_222650_TOF_Pyruvic acid (1MEOX) (1TMS | Pyruvate | Acid (Oxo) | C3H4O3 | 174 | Tomato, Arabidopsis, potato, grapevine and others | | A |
| 8 | 2 | 271580 | 271806 | 0,1 | M000030_A122001-101_METB_271580_TOF_Valine, DL- (2TMS) | Valine | Acid (Amino) | C5H11NO2 | 144 | Tomato, Arabidopsis, potato, grapevine and others | | A |
| 9 | 3 | 291783 | 291776 | -0,03 | M000053_A129003-101_METB_291783_TOF_Glycerol (3TMS) | Glycerol | Polyol | C3H8O3 | 205 | Tomato, Arabidopsis, potato, grapevine and others | | A |
| 10 | 4 | 319193 | 319036 | -0,03 | M000017_A132002-101_METB_319193_TOF_Isoleucine, L- (2TMS) | Isoleucine | Acid (Amino) | C6H13NO2 | 158 | Tomato, Arabidopsis, potato, grapevine and others | | A |
| 11 | 5 | 325180 | 324856 | -0,1 | M000031_A133001-101_METB_325180_TOF_Glycine (3TMS) | Glycine | Acid (Amino) | C2H5NO2 | 174 | Tomato, Arabidopsis, potato, grapevine and others | | A |
| 12 | 6 | 333520 | 332443 | -0,37 | M000075_A129001-101_METB_333520_TOF_Phosphoric acid (3TMS) | Phosphorate | Phosphate | H3O4P | 299 | Tomato, Arabidopsis, potato, grapevine and others | | A |
| 13 | 7 | 338693 | 338620 | -0,04 | M000029_A132003-101_METB_338693_TOF_Proline, L- (2TMS) | Proline | Amide | CH4N2O | 142 | Tomato, Arabidopsis, potato, grapevine and others | | A |
| 14 | 8 | 340357 | 339818 | -0,2 | M000364_A127002-101_METB_340357_TOF_Urea (2TMS) | Urea | Amide | CH4N2O | 189 | Tomato, Arabidopsis, potato, grapevine and others | | A |
| 15 | 9 | 342530 | 342246 | -0,07 | M000114_A133002-101_METB_342530_TOF_Butyric acid, 4-amino- (2TMS | 4-Aminobutanoic Acid | Acid (Amino) | C4H9NO2 | 103 | Tomato, Arabidopsis, potato, grapevine and others | | A |
| 16 | 10 | 345050 | 344205 | -0,23 | M000073_A135003-101_METB_345050_TOF_Glyceric acid, DL- (3TMS) | Glycerate | Acid | C3H6O4 | 292 | Tomato, Arabidopsis, potato, grapevine and others | | A |
| 17 | 11 | 354060 | 353867 | -0,06 | M000026_A138002-101_METB_354060_TOF_Alanine, DL- (3TMS) | Alanine | Acid (Amino) | C3H7NO2 | 188 | Tomato, Arabidopsis, potato, grapevine and others | | A |
| 18 | 12 | 357523 | 357021 | -0,13 | M000015_A138001-101_METB_357523_TOF_Serine, DL- (3TMS) | Serine | Acid (Amino) | C3H7NO3 | 204 | Tomato, Arabidopsis, potato, grapevine and others | | A |
| 19 | 13 | 365427 | 364803 | -0,13 | M000074_A134001-101_METB_365427_TOF_Succinic acid (2TMS) | Succinate | Acid | C4H6O4 | 247 | Tomato, Arabidopsis, potato, grapevine and others | | A |
| # | 14 | 368365 | 367319 | -0,3 | M000016_A140001-101_METB_368365_TOF_Threonine, DL- (3TMS) | Threonine | Acid (Amino) | C4H9NO3 | 117 | Tomato, Arabidopsis, potato, grapevine and others | | A |
| 21 | 15 | 371255 | 370722 | -0,15 | M000067_A137001-101_METB_371255_TOF_Fumaric acid (2TMS) | Fumarate | Acid (Dicarboxylic) | C4H4O4 | 245 | Tomato, Arabidopsis, potato, grapevine and others | | A |
| # | 16 | 386210 | 385684 | -0,14 | M000457_A133004-101_METB_386210_TOF_Nicotinic acid (1TMS) | Nicotinate | Acid | C6H5NO2 | 180 | Tomato, Arabidopsis, potato, grapevine and others | | A |
| # | 17 | 394250 | 393982 | -0,06 | M000027_A144001-101_METB_394250_TOF_Alanine, beta- (3TMS) | beta-Alanine | Acid (Amino) | C3H7NO2 | 248 | Tomato, Arabidopsis, potato, grapevine and others | | A |
| # | 18 | 404940 | 405841 | 0,21 | M000019_A146001-101_METB_404940_TOF_Homoserine, DL- (3TMS) | Homoserine | Acid (Amino) | C4H9NO3 | 119 | Tomato, Arabidopsis, potato, grapevine and others | | A |
| 25 | 19 | 408455 | 408118 | -0,06 | M000054_A150002-101_METB_408455_TOF_Erythritol (4TMS) | Erythritol | Polyol | C4H10O4 | 205 | Tomato, Arabidopsis, potato, grapevine and others | | A |
| # | 20 | 440995 | 440474 | -0,11 | M000065_A149001-101_METB_440995_TOF_Malic acid, DL- (3TMS) | Malate | Acid | C4H6O5 | 233 | Tomato, Arabidopsis, potato, grapevine and others | | A |
| 27 | 21 | 457283 | 456775 | -0,03 | M000033_A152002-101_METB_457283_TOF_Aspartic acid, L- (3TMS) | Aspartate | Acid (Amino) | C4H7NO4 | 232 | Tomato, Arabidopsis, potato, grapevine and others | | A |
| # | 22 | 474325 | 473799 | -0,11 | M000018_A152001-101_METB_474325_TOF_Methionine, DL- (2TMS) | Methionine | Acid (Amino) | C5H11NO2S | 176 | Tomato, Arabidopsis, potato, grapevine and others | | A |
| # | 23 | 492650 | 492221 | -0,08 | M000032_A174008-101_MST_492650_TOF_Glutamine, DL- (4TMS) | Glutamine | Acid (Amino) | C5H10N2O3 | 227 | Tomato, Arabidopsis, potato, grapevine and others | | A |
| # | 24 | 498150 | 497775 | -0,04 | M000579_A166001-101_METB_498150_TOF_Xylose, D- (1MEOX) (4TMS) | Xylose | Sugar (Pentose, aldos | C5H10O5 | 160 | Tomato, Arabidopsis, potato, grapevine and others | | A |
| 31 | 25 | 507780 | 507375 | -0,09 | M000036_A163001-101_METB_507780_TOF_Glutamic acid, DL- (3TMS) | Glutamate | Acid (Amino) | C5H9NO4 | 246 | Tomato, Arabidopsis, potato, grapevine and others | | A |
| # | 26 | 517180 | 517566 | 0,1 | M000186_A175002-101_METB-METB_517180_TOF_Putrescine (4TMS) | Putrescine | Amine | C4H12N2 | 174 | Tomato, Arabidopsis, potato, grapevine and others | | A |
| # | 27 | 523847 | 522791 | -0,19 | M000571_A158004-101_METB_523847_TOF_Glutaric acid, 2-oxo- (1MEOX) | 2-oxo-Glutarate | Acid (Oxo) | C5H6O5 | 198 | Tomato, Arabidopsis, potato, grapevine and others | | A |
| # | 28 | 531145 | 530508 | -0,11 | M000011_A164001-101_METB_531145_TOF_Phenylalanine, DL- (2TMS) | Phenylalanine | Acid (Amino) | C9H11NO2 | 218 | Tomato, Arabidopsis, potato, grapevine and others | | A |
| 35 | 29 | 550100 | 549546 | -0,1 | M000013_A168001-101_METB_550100_TOF_Asparagine, DL- (3TMS) | Asparagine | Acid (Amino) | C4H8N2O3 | 116 | Tomato, Arabidopsis, potato, grapevine and others | | A |
| # | 30 | 570427 | 570449 | 0,03 | M000028_A182002-101_METB-METB_570427_TOF_Ornithine, DL- (4TMS | Ornithine | Acid (Amino) | C5H12N2O2 | 142 | Tomato, Arabidopsis, potato, grapevine and others | | A |
| 37 | 31 | 574230 | 573233 | -0,18 | M000328_A177002-101_METB_574230_TOF_Glycerol-3-phosphate, DL- (4 | Glycerol-3-P | Alcohol (Phosphate) | C3H9O6P | 299 | Tomato, Arabidopsis, potato, grapevine and others | | A |
| # | 48 | 579380 | 576444 | -0,5 | M000606_A187002-101_METB_579380_TOF_Fructose, D- (1MEOX) (5TMS | Fructose | Sugar (Hexose, ketose | C6H12O6 | 217 | Tomato, Arabidopsis, potato, grapevine and others | | A |
| # | 49 | 590400 | 587560 | -0,44 | M000040_A189002-101_METB_590400_TOF_Glucose, D- (1MEOX) (5TMS | Glucose | Sugar (Hexose, aldose | C6H12O6 | 160 | Tomato, Arabidopsis, potato, grapevine and others | | A |
| # | 50 | 592883 | 592883 | 0 | M000069_A182004-101_METB-METB_592883_TOF_Citric acid (4TMS) | Citrate | Acid | C6H8O7 | 273 | Tomato, Arabidopsis, potato, grapevine and others | | A |
| 41 | 32 | 615467 | 616388 | 0,13 | M000014_A192003-101_METB_615467_TOF_Lysine, L- (4TMS) | Lysine | Acid (Amino) | C6H14N2O2 | 156 | Tomato, Arabidopsis, potato, grapevine and others | | A |
| # | 33 | 618870 | 618615 | -0,13 | M000639_A189003-101_METB_618870_TOF_Galactonic acid-1,4-lactone, D | D-Galactono-1,4-lactone | Acid (Hexonic) | C6H12O7 | 178 | Tomato, Arabidopsis, potato, grapevine and others | | A |
| # | 34 | 624990 | 624222 | -0,06 | M000082_A185002-101_METB-METB_624990_TOF_Dehydroascorbic acid | Dehydroascorbate | Acid | C6H6O6 | 316 | Tomato, Arabidopsis, potato, grapevine and others | | A |
| # | 35 | 632880 | 632250 | -0,08 | M000690_A196003-101_METB_632880_TOF_Galacturonic acid, D- (1MEO | Galacturonate | Acid (Hexuronic) | C6H10O7 | 194 | Tomato, Arabidopsis, potato, grapevine and others | | A |
| 45 | 36 | 653910 | 653994 | 0,04 | M000060_A209002-101_METB_653910_TOF_Inositol, myo- (6TMS) | myo-Inositol | Polyol | C6H12O6 | 305 | Tomato, Arabidopsis, potato, grapevine and others | | A |
| # | 37 | 658337 | 657691 | -0,09 | M000035_A194002-101_METB_658337_TOF_Tyrosine, DL- (3TMS) | Tyrosine | Acid (Amino) | C9H11NO3 | 218 | Tomato, Arabidopsis, potato, grapevine and others | | A |
| 47 | 38 | 767060 | 765960 | -0,15 | M000510_A232002-101_METB_767060_TOF_Fructose-6-phosphate (1MEO | Fructose-6-phosphate | Sugar (Phosphate) | C6H13O9P | 260 | Tomato, Arabidopsis, potato, grapevine and others | | A |
| # | 39 | 781350 | 778808 | -0,35 | M000513_A235002-101_METB-METB_781350_TOF_Glucose-6-phosphate ( | Glucose-6-phosphate | Sugar (Phosphate) | C6H13O9P | 260 | Tomato, Arabidopsis, potato, grapevine and others | | A |
| # | 40 | 790560 | 790597 | 0 | M000012_A223001-101_METB_790560_TOF_Tryptophan, L- (3TMS) | Tryptophan | Acid (Amino) | C11H12N2O2 | 204 | Tomato, Arabidopsis, potato, grapevine and others | | A |
| 50 | 47 | 840783 | 837519 | -0,37 | M000044_A264001-101_METB_840783_TOF_Sucrose, D- (8TMS) | Sucrose | Sugar (Disaccharide) | C12H22O11 | 361 | Tomato, Arabidopsis, potato, grapevine and others | | A |
| 51 | 41 | 870355 | 868638 | -0,2 | M000048_A274001-101_METB_870355_TOF_Maltose, D- (1MEOX) (8TMS | Maltose | Sugar (Disaccharide) | C12H22O11 | 204 | Tomato, Arabidopsis, potato, grapevine and others | | A |
| 52 | 42 | 876240 | 874646 | -0,17 | M000671_A274002-101_METB_876240_TOF_Trehalose, alpha,alpha'-, D- (8 | Trehalose | Sugar (Disaccharide) | C12H22O11 | 191 | Tomato, Arabidopsis, potato, grapevine and others | | A |
| 53 | 43 | 940280 | 938445 | -0,19 | M000673_A299002-101_METB_940280_EIROE_Galactinol (9TMS) | Galactinol | Polyol | C12H22O11 | 204 | Tomato, Arabidopsis, potato, grapevine and others | | A |
| 54 | 44 | 996820 | 995772 | -0,11 | M000644_A299001-101_METB_996820_TOF_Quinic acid, 3-caffeoyl-, cis- | Quinic acid, 3-caffeoyl-, cis- | Conjugate (Phenylprop | C16H18O9 | 787 | Tomato, Arabidopsis, potato, grapevine and others | | A |
| 55 | 45 | 1025600 | 1024785 | -0,08 | M000003_A311001-101_METB_1025600_TOF_Quinic acid, 3-caffeoyl-, trans | Quinic acid, 3-caffeoyl-, trans- | Conjugate (Phenylprop | C16H18O9 | 787 | Tomato, Arabidopsis, potato, grapevine and others | | A |
| 56 | 46 | 1033367 | 1031719 | -0,16 | M000049_A337002-101_METB_1033367_TOF_Raffinose (11TMS) | Raffinose | Sugar (Trisaccharide) | C18H32O16 | 361 | Tomato, Arabidopsis, potato, grapevine and others | | A |
| 57 | 47 | 1054900 | 1054424 | -0,05 | M000284_A307003-101_METB_1054900_TOF_Adenosine-5-monophospha | Adenosine-5-monophosphate | Nucleotide (Monopho | C10H14N5O7P | 347 | Tomato, Arabidopsis, potato, grapevine and others | | A |

**Table S3 Parameters used for peak annotation of all detected metabolites in WT-MicroTom and *aox1a* mutant fruits.**

**Supplemental Table 3.** Overview of the metabolite reporting list.

**GC-TOF-MS metabolites**

| | A | B | C | D | E | F | G | H | I | J | K | L |
|---|---|---|---|---|---|---|---|---|---|---|---|---|
| 1 | **Experiment title:** | | | Metabolite profiles of tomato wild-type and *aox1a* mutants during ripening | | Peak/compound no.- number referenced back to the main text | | | Mass to charge ratio (m/z) | | | |
| 2 | **Organism/Plant species:** | | | Solanum lycopersicum | | Ret . Time- Time expected, Tag Time Index and Time deviation | | | (S)- identification confirmed by a standard compound | | | |
| 3 | **Organ/tissue:** | | | Fruit Pericarp | | Putative Name- putative identification of the metabolite/derivativ | | | I, II, III- different isomers | | | |
| 4 | **Analytical tool:** | | | GC-TOF-MS | | Corresponding metabolite name in literature | | | Identification level (A; B; C; D)- (A) standard or NMR; (B) MS/MS; (C) MS$^E$; (D) MS onl | | | |
| 5 | | | | | | Mol. Formula- molecular formula of the metabolite or its FA adduct; | | | | | | |
| 6 | **Peak/ Compound** | **Time Expected** | **Tag Time Index** | **Time Deviation** | **Putative metabolite name (Derivative)** | **Corresponding Metabolite in Literature** | **Metabolite Class** | **Mol formula** | **Mass to charge ratio** | **Species detected before** | **Ref** | **Identification level (A-D)** |
| 7 | 1 | [illegible] | [illegible] | [illegible] | M000071_A104002-101_METB_222650_TOF_Pyruvic acid (1MEOX) (1TMS) | Pyruvate | Acid (Oxo) | C3H4O3 | 174 | Tomato, Arabidopsis, potato, grapevine and others | | A |
| 8 | 2 | [illegible] | [illegible] | [illegible] | M000030_A122001-101_METB_271580_TOF_Valine, DL- (2TMS) | Valine | Acid (Amino) | C5H11NO2 | 144 | Tomato, Arabidopsis, potato, grapevine and others | | A |
| 9 | 3 | [illegible] | [illegible] | [illegible] | M000053_A129003-101_METB_291783_TOF_Glycerol (3TMS) | Glycerol | Polyol | C3H8O3 | 205 | Tomato, Arabidopsis, potato, grapevine and others | | A |
| 10 | 4 | [illegible] | [illegible] | [illegible] | M000017_A132002-101_METB_319193_TOF_Isoleucine, L- (2TMS) | Isoleucine | Acid (Amino) | C6H13NO2 | 158 | Tomato, Arabidopsis, potato, grapevine and others | | A |
| 11 | 5 | [illegible] | [illegible] | [illegible] | M000031_A133001-101_METB_325180_TOF_Glycine (3TMS) | Glycine | Acid (Amino) | C2H5NO2 | 174 | Tomato, Arabidopsis, potato, grapevine and others | | A |
| 12 | 6 | [illegible] | [illegible] | [illegible] | M000075_A129001-101_METB_333520_TOF_Phosphoric acid (3TMS) | Phosphorate | Phosphate | H3O4P | 225 | Tomato, Arabidopsis, potato, grapevine and others | | A |
| 13 | 7 | [illegible] | [illegible] | [illegible] | M000029_A132003-101_METB_338693_TOF_Proline, L- (2TMS) | Proline | Amide | CH4N2O | 142 | Tomato, Arabidopsis, potato, grapevine and others | | A |
| 14 | 8 | [illegible] | [illegible] | [illegible] | M000364_A127002-101_METB_340357_TOF_Urea (2TMS) | Urea | Amide | CH4N2O | 171 | Tomato, Arabidopsis, potato, grapevine and others | | A |
| 15 | 9 | [illegible] | [illegible] | [illegible] | M000073_A135003-101_METB_345050_TOF_Glyceric acid, DL- (3TMS) | Glycerate | Acid | C3H6O4 | 189 | Tomato, Arabidopsis, potato, grapevine and others | | A |
| 16 | 10 | [illegible] | [illegible] | [illegible] | M000015_A138001-101_METB_357523_TOF_Serine, DL- (3TMS) | Serine | Acid (Amino) | C3H7NO3 | 204 | Tomato, Arabidopsis, potato, grapevine and others | | A |
| 17 | 11 | [illegible] | [illegible] | [illegible] | M000074_A134001-101_METB_365427_TOF_Succinic acid (2TMS) | Succinate | Acid | C4H6O4 | 247 | Tomato, Arabidopsis, potato, grapevine and others | | A |
| 18 | 12 | [illegible] | [illegible] | [illegible] | M000016_A140001-101_METB_368365_TOF_Threonine, DL- (3TMS) | Threonine | Acid (Amino) | C4H9NO3 | 219 | Tomato, Arabidopsis, potato, grapevine and others | | A |
| 19 | 13 | [illegible] | [illegible] | [illegible] | M000067_A137001-101_METB_371255_TOF_Fumaric acid (2TMS) | Fumarate | Acid (Dicarboxylic) | C4H4O4 | 245 | Tomato, Arabidopsis, potato, grapevine and others | | A |
| # | 14 | [illegible] | [illegible] | [illegible] | M000457_A133004-101_METB_386210_TOF_Nicotinic acid (1TMS) | Nicotinate | Acid | C6H5NO2 | 180 | Tomato, Arabidopsis, potato, grapevine and others | | A |
| 21 | 15 | [illegible] | [illegible] | [illegible] | M000027_A144001-101_METB_394250_TOF_Alanine, beta- (3TMS) | beta-Alanine | Acid (Amino) | C3H7NO2 | 174 | Tomato, Arabidopsis, potato, grapevine and others | | A |
| # | 16 | [illegible] | [illegible] | [illegible] | M000019_A146001-101_METB_404940_TOF_Homoserine, DL- (3TMS) | Homoserine | Acid (Amino) | C4H9NO3 | 128 | Tomato, Arabidopsis, potato, grapevine and others | | A |
| # | 17 | [illegible] | [illegible] | [illegible] | M000054_A150002-101_METB_408455_TOF_Erythritol (4TMS) | Erythritol | Polyol | C4H10O4 | 217 | Tomato, Arabidopsis, potato, grapevine and others | | A |
| # | 18 | [illegible] | [illegible] | [illegible] | M000065_A149001-101_METB_440995_TOF_Malic acid, DL- (3TMS) | Malate | Acid | C4H6O5 | 233 | Tomato, Arabidopsis, potato, grapevine and others | | A |
| # | 19 | [illegible] | [illegible] | [illegible] | M000114_A133002-101_METB_342530_TOF_Butyric acid, 4-amino- (2TMS) | 4-Aminobutanoic Acid | Acid (Amino) | C4H9NO2 | 246 | Tomato, Arabidopsis, potato, grapevine and others | | A |
| # | 20 | [illegible] | [illegible] | [illegible] | M000033_A152002-101_METB_457283_TOF_Aspartic acid, L- (3TMS) | Aspartate | Acid (Amino) | C4H7NO4 | 232 | Tomato, Arabidopsis, potato, grapevine and others | | A |
| # | 21 | [illegible] | [illegible] | [illegible] | M000018_A152001-101_METB_474325_TOF_Methionine, DL- (2TMS) | Methionine | Acid (Amino) | C5H11NO2S | 176 | Tomato, Arabidopsis, potato, grapevine and others | | A |
| # | 22 | [illegible] | [illegible] | [illegible] | M000013_A168001-101_METB_550100_TOF_Asparagine, DL- (3TMS) | Asparagine | Acid (Amino) | C4H8N2O3 | [illegible] | Tomato, Arabidopsis, potato, grapevine and others | | A |
| # | 23 | [illegible] | [illegible] | [illegible] | M000028_A182002-101_METB-METB_570427_TOF_Ornithine, DL- (4TMS) | Ornithine | Acid (Amino) | C5H12N2O2 | 142 | Tomato, Arabidopsis, potato, grapevine and others | | A |
| # | 24 | [illegible] | [illegible] | [illegible] | M000032_A174008-101_MST_492650_TOF_Glutamine, DL- (4TMS) | Glutamine | Acid (Amino) | C5H10N2O3 | 227 | Tomato, Arabidopsis, potato, grapevine and others | | A |
| 31 | 25 | [illegible] | [illegible] | [illegible] | M000579_A166001-101_METB_498150_TOF_Xylose, D- (1MEOX) (4TMS) | Xylose | Sugar (Pentose, aldose | C5H10O5 | 217 | Tomato, Arabidopsis, potato, grapevine and others | | A |
| # | 26 | [illegible] | [illegible] | [illegible] | M000036_A163001-101_METB_507780_TOF_Glutamic acid, DL- (3TMS) | Glutamate | Acid (Amino) | C5H9NO4 | 246 | Tomato, Arabidopsis, potato, grapevine and others | | A |
| # | 27 | [illegible] | [illegible] | [illegible] | M000590_A172002-101_METB_515530_TOF_Rhamnose, DL- (1MEOX) (4TMS) | Rhamnose | Sugar (Hexose, deoxy) | C6H12O5 | 117 | Tomato, Arabidopsis, potato, grapevine and others | | A |
| # | 28 | [illegible] | [illegible] | [illegible] | M000186_A175002-101_METB-METB_517180_TOF_Putrescine (4TMS) | Putrescine | Amine | C4H12N2 | 174 | Tomato, Arabidopsis, potato, grapevine and others | | A |
| # | 29 | [illegible] | [illegible] | [illegible] | M000591_A173002-101_METB_522140_TOF_Fucose, DL- (1MEOX) (4TMS) | Fucose | Sugar (Hexose, deoxy) | C6H12O5 | 117 | Tomato, Arabidopsis, potato, grapevine and others | | A |
| # | 30 | [illegible] | [illegible] | [illegible] | M000571_A158004-101_METB_523847_TOF_Glutaric acid, 2-oxo- (1MEOX) (2TMS) | 2-oxo-Glutarate | Acid (Oxo) | C5H6O5 | [illegible] | Tomato, Arabidopsis, potato, grapevine and others | | A |
| # | 31 | [illegible] | [illegible] | [illegible] | M000011_A164001-101_METB_531145_TOF_Phenylalanine, DL- (2TMS) | Phenylalanine | Acid (Amino) | C9H11NO2 | 192 | Tomato, Arabidopsis, potato, grapevine and others | | A |
| # | 32 | [illegible] | [illegible] | [illegible] | M000592_A172006-101_METB_553540_TOF_Adipic acid, 2-amino-, DL- (3TMS) | 2-amino-Adipic acid | Acid (Amino) | C6H11NO4 | 217 | Tomato, Arabidopsis, potato, grapevine and others | | A |
| # | 33 | [illegible] | [illegible] | [illegible] | M000360_A181006-101_METB_563070_TOF_Calystegine B2 (1MEOX) (4TMS) | Calystegine B2 | Calystegine | C7H13NO4 | 257 | Tomato, Arabidopsis, potato, grapevine and others | | A |
| # | 34 | [illegible] | [illegible] | [illegible] | M000328_A177002-101_METB_574230_TOF_Glycerol-3-phosphate, DL- (4TMS) | Glycerol-3-P | Alcohol (Phosphate) | C3H9O6P | 299 | Tomato, Arabidopsis, potato, grapevine and others | | A |
| 41 | 35 | [illegible] | [illegible] | [illegible] | M000007_A185001-101_METB_577930_TOF_Quinic acid, D(-)- (5TMS) | Quinate | Acid | C7H12O6 | 255 | Tomato, Arabidopsis, potato, grapevine and others | | A |
| # | 36 | [illegible] | [illegible] | [illegible] | M000606_A187002-101_METB_579380_TOF_Fructose, D- (1MEOX) (5TMS) | Fructose | Sugar (Hexose, ketose | C6H12O6 | 307 | Tomato, Arabidopsis, potato, grapevine and others | | A |
| # | 37 | [illegible] | [illegible] | [illegible] | M000069_A182004-101_METB-METB_592883_TOF_Citric acid (4TMS) | Citrate | Acid | C6H8O7 | 273 | Tomato, Arabidopsis, potato, grapevine and others | | A |
| # | 38 | [illegible] | [illegible] | [illegible] | M000608_A182003-101_METB-METB_597845_TOF_Isocitric acid, DL- (4TMS) | Isocitrate | Acid | C6H8O7 | 465 | Tomato, Arabidopsis, potato, grapevine and others | | A |
| # | 39 | [illegible] | [illegible] | [illegible] | M000040_A189002-101_METB_590400_TOF_Glucose, D- (1MEOX) (5TMS) | Glucose | Sugar (Hexose, aldose | C6H12O6 | 160 | Tomato, Arabidopsis, potato, grapevine and others | | A |
| # | 40 | [illegible] | [illegible] | [illegible] | M000639_A189003-101_METB_618870_TOF_Galactonic acid-1,4-lactone, D(-)- (4TMS | D-Galactono-1,4-lactone | Acid (Hexonic) | C6H12O7 | 217 | Tomato, Arabidopsis, potato, grapevine and others | | A |
| # | 41 | [illegible] | [illegible] | [illegible] | M000014_A192003-101_METB_615467_TOF_Lysine, L- (4TMS) | Lysine | Acid (Amino) | C6H14N2O2 | 305 | Tomato, Arabidopsis, potato, grapevine and others | | A |
| # | 42 | [illegible] | [illegible] | [illegible] | M000639_A189003-101_METB_618870_TOF_Galactonic acid-1,4-lactone, D(-)- (4TMS | Galactonic acid-1,4-lactone | Acid (polyhydroxy) | C6H10O6 | 173 | Tomato, Arabidopsis, potato, grapevine and others | | A |
| # | 43 | [illegible] | [illegible] | [illegible] | M000082_A185002-101_METB-METB_624990_TOF_Dehydroascorbic acid dimer (TM | Dehydroascorbate | Acid | C6H6O6 | 316 | Tomato, Arabidopsis, potato, grapevine and others | | A |
| # | 44 | [illegible] | [illegible] | [illegible] | M000081_A191004-101_METB_638360_TOF_Tyramine (3TMS) | Tyramine | Amine | C8H11NO | 174 | Tomato, Arabidopsis, potato, grapevine and others | | A |
| 51 | 45 | [illegible] | [illegible] | [illegible] | M000060_A209002-101_METB_653910_TOF_Inositol, myo- (6TMS) | myo-Inositol | Polyol | C6H12O6 | 191 | Tomato, Arabidopsis, potato, grapevine and others | | A |
| # | 46 | [illegible] | [illegible] | [illegible] | M000035_A194002-101_METB_658337_TOF_Tyrosine, DL- (3TMS) | Tyrosine | Acid (Amino) | C9H11NO3 | 218 | Tomato, Arabidopsis, potato, grapevine and others | | A |
| # | 47 | [illegible] | [illegible] | [illegible] | M000643_A194009-101_METB_660120_TOF_Glucaric acid-1,4-lactone (4TMS) | Glucarate | Acid (polyhydroxy) | C6H8O7 | 217 | Tomato, Arabidopsis, potato, grapevine and others | | A |
| # | 48 | [illegible] | [illegible] | [illegible] | M000039_A192006-101_METB_679450_TOF_Histidine, L- (3TMS) | Histidine | Acid (Amino) | C6H9N3O2 | 154 | Tomato, Arabidopsis, potato, grapevine and others | | A |
| 55 | 49 | [illegible] | [illegible] | [illegible] | M000106_A220002-101_METB_728760_TOF_Spermidine (4TMS) | Spermidine | Amine | C7H19N3 | 144 | Tomato, Arabidopsis, potato, grapevine and others | | A |
| # | 50 | [illegible] | [illegible] | [illegible] | M000510_A232002-101_METB_767060_TOF_Fructose-6-phosphate (1MEOX) (6TMS) | Fructose-6-phosphate | Sugar (Phosphate) | C6H13O9P | 315 | Tomato, Arabidopsis, potato, grapevine and others | | A |
| 57 | 51 | [illegible] | [illegible] | [illegible] | M000513_A235002-101_METB-METB_781350_TOF_Glucose-6-phosphate (1MEOX) (6 | Glucose-6-phosphate | Sugar (Phosphate) | C6H13O9P | 87 | Tomato, Arabidopsis, potato, grapevine and others | | A |
| # | 52 | [illegible] | [illegible] | [illegible] | M000012_A223001-101_METB_790560_TOF_Tryptophan, L- (3TMS) | Tryptophan | Acid (Amino) | C11H12N2O2 | 202 | Tomato, Arabidopsis, potato, grapevine and others | | A |
| # | 53 | [illegible] | [illegible] | [illegible] | M000044_A264001-101_METB_840783_TOF_Sucrose, D- (8TMS) | Sucrose | Sugar (Disaccharide) | C12H22O11 | 361 | Tomato, Arabidopsis, potato, grapevine and others | | A |
| # | 54 | [illegible] | [illegible] | [illegible] | M000671_A274002-101_METB_876240_TOF_Trehalose, alpha,alpha'-, D- (8TMS) | Trehalose | Sugar (Disaccharide) | C12H22O11 | 191 | Tomato, Arabidopsis, potato, grapevine and others | | A |
| 61 | 55 | [illegible] | [illegible] | [illegible] | M000048_A274001-101_METB_870355_TOF_Maltose, D- (1MEOX) (8TMS) | Maltose | Sugar (Disaccharide) | C12H22O11 | 361 | Tomato, Arabidopsis, potato, grapevine and others | | A |
| # | 56 | [illegible] | [illegible] | [illegible] | M000107_A287001-101_METB_909948_TOF_Isomaltose (1MEOX) (8TMS) | Isomaltose | Sugar (Disaccharide) | C12H22O11 | 204 | Tomato, Arabidopsis, potato, grapevine and others | | A |

**Table S4 Relative metabolite levels detected by GC-MS in WT-Ailsa Craig tomato fruits.** Values were determined after normalization by the mean levels of WT pericarp samples at mature green (MG) stage. Levels correspond to the mean ± SE of n=4-6 independent replicates. Asterisks denote significant differences (P < 0.05) between MG and other ripening stages (BR, OR, R). 'n.d.' denotes 'not detected'.

| | MG | BR | OR | RR |
|---|---|---|---|---|
| Pyruvate | 1 ± 0,13 | 1,87 ± 0,2* | 1,52 ± 0,12* | 1,36 ± 0,14 |
| Valine | 1 ± 0,12 | 0,22 ± 0,03* | 0,14 ± 0,06* | 0,39 ± 0,11* |
| Isoleucine | 1 ± 0,08 | 0,5 ± 0,07* | 0,32 ± 0,1* | 0,7 ± 0,18 |
| Glycine | 1 ± 0,25 | 0,35 ± 0,05* | 0,27 ± 0,08* | 0,73 ± 0,22 |
| Phosphoric acid | 1 ± 0,08 | 0,91 ± 0,04 | 0,92 ± 0,04 | 0,99 ± 0,07 |
| Proline | 1 ± 0,18 | 0,49 ± 0,11* | 0,47 ± 0,11* | 3,3 ± 1,09 |
| Urea | 1 ± 0,1 | 0,74 ± 0,14 | 0,64 ± 0,12* | 0,8 ± 0,27 |
| GABA | 1 ± 0,16 | 0,64 ± 0,14 | 0,44 ± 0,12* | 0,34 ± 0,12* |
| Glycerate | 1 ± 0,06 | 0,65 ± 0,08* | 0,52 ± 0,14* | 0,5 ± 0,09* |
| Alanine | 1 ± 0,14 | 0,41 ± 0,04* | 0,7 ± 0,2 | 3,38 ± 1,34 |
| Serine | 1 ± 0,11 | 0,83 ± 0,21 | 0,57 ± 0,13* | 0,85 ± 0,17 |
| Succinante | 1 ± 0,06 | 2,37 ± 0,18* | 1,98 ± 0,34* | 2,92 ± 0,59* |
| Threonine | 1 ± 0,09 | 0,66 ± 0,11* | 0,55 ± 0,17* | 0,66 ± 0,16 |
| Fumarate | 1 ± 0,11 | 1,51 ± 0,27 | 1,4 ± 0,12* | 1,84 ± 0,31* |
| Nicotinate | 1 ± 0,15 | 0,81 ± 0,08 | 0,82 ± 0,12 | 0,76 ± 0,08 |
| Alanine, beta | 1 ± 0,18 | 0,41 ± 0,09* | 0,22 ± 0,06* | 0,27 ± 0,1* |
| Homoserine | 1 ± 0,15 | 0,97 ± 0,1 | 1,08 ± 0,2 | 1,15 ± 0,26 |
| Erythritol | 1 ± 0,1 | 1,01 ± 0,09 | 1,38 ± 0,15 | 1,78 ± 0,32* |
| Methionine | 1 ± 0,19 | 1,75 ± 0,21* | 4,11 ± 0,55* | 3,08 ± 0,59* |
| Glutamine | 1 ± 0,17 | 1,22 ± 0,23 | 0,97 ± 0,33 | 0,56 ± 0,22 |
| Xylose | 1 ± 0,07 | 1,45 ± 0,1* | 1,66 ± 0,17* | 1,56 ± 0,07* |
| Putrescine | 1 ± 0,17 | 0,73 ± 0,21 | 0,75 ± 0,26 | 0,65 ± 0,17 |
| 2-oxo-Glutarate | 1 ± 0,1 | 3,35 ± 0,11* | 5,14 ± 0,53* | 9,6 ± 2,07* |
| Phenylalanine | 1 ± 0,1 | 0,63 ± 0,2 | 0,41 ± 0,15* | 0,41 ± 0,09* |
| Asparagine | 1 ± 0,26 | 0,99 ± 0,18 | 1,01 ± 0,35 | 0,63 ± 0,18 |
| Ornithine | 1 ± 0,14 | 0,88 ± 0,2 | 1,04 ± 0,29 | 0,89 ± 0,21 |
| Lysine | 1 ± 0,11 | 0,88 ± 0,14 | 0,73 ± 0,21 | 0,83 ± 0,17 |
| Galactonate 1,4-lactone | 1 ± 0,12 | 0,89 ± 0,08 | 1,31 ± 0,17 | 1,43 ± 0,2 |
| Dehydroascorbate | 1 ± 0,15 | 2,29 ± 0,08* | 2,3 ± 0,1* | 2,91 ± 0,37* |
| Galacturonate | 1 ± 0,06 | 3,26 ± 0,35* | 45,69 ± 12,35* | 46,49 ± 8,01* |
| myo-Inositol | 1 ± 0,08 | 0,85 ± 0,1 | 0,77 ± 0,07 | 1,14 ± 0,05 |
| Tyrosine | 1 ± 0,2 | 0,4 ± 0,1* | 0,32 ± 0,11* | 0,5 ± 0,21 |
| Fructose-6-P | 1 ± 0,12 | 0,88 ± 0,06 | 1,35 ± 0,14 | 1,67 ± 0,12* |
| Tryptophan | 1 ± 0,15 | 0,48 ± 0,09* | 0,67 ± 0,25 | 1,14 ± 0,19 |
| Maltose | 1 ± 0,15 | 0,79 ± 0,08 | 1,11 ± 0,17 | 1,79 ± 0,25* |
| Trehalose | 1 ± 0,18 | 1,34 ± 0,07 | 1,6 ± 0,27 | 2,43 ± 0,19* |
| Galactinol | 1 ± 0,39 | n.d. | 1,65 ± 1,19 | 0,84 ± 0,1 |
| 3-caffeoyl, cis-Quinate | 1 ± 0,23 | 1,04 ± 0,26 | 0,62 ± 0,11 | 0,83 ± 0,21 |
| 3-caffeoyl, trans-Quinate | 1 ± 0,28 | 1,47 ± 0,24 | 1,4 ± 0,62 | 1,02 ± 0,35 |
| Raffinose | 1 ± 0,19 | 0,46 ± 0,06* | 0,66 ± 0,38 | 0,47 ± 0,12 |
| Adenosine-5-monophosphate | 1 ± 0,18 | 2,29 ± 0,48* | 9,28 ± 1,45* | 14,02 ± 4,26 |
| Malate | 1 ± 0,16 | 1,16 ± 0,16 | 0,9 ± 0,11 | 0,4 ± 0,08* |
| Aspartate | 1 ± 0,1 | 1,79 ± 0,33* | 2,3 ± 0,64 | 1,5 ± 0,33 |
| Glutamate | 1 ± 0,21 | 1,28 ± 0,2 | 1,9 ± 0,58 | 1,53 ± 0,47 |
| Fructose | 1 ± 0,05 | 1,09 ± 0,05 | 1,15 ± 0,05 | 1,35 ± 0,05* |
| Glucose | 1 ± 0,05 | 1,03 ± 0,04 | 1,01 ± 0,08 | 1,35 ± 0,09* |
| Citrate | 1 ± 0,16 | 1,74 ± 0,15* | 2,2 ± 0,11* | 2,49 ± 0,14* |
| Sucrose | 1 ± 0,1 | 0,38 ± 0,05* | 0,26 ± 0,05* | 0,39 ± 0,06* |

**Table S5 Relative metabolite levels detected by GC-MS in *ghost* tomato fruits.** Values were determined after normalization by the mean levels of WT pericarp samples at mature white (MW) stage. Levels correspond to the mean ± SE of n=4 independent replicates. Asterisks denote significant differences (P < 0.05) between MG and other ripening stages (BR and Y). 'n.d.' denotes 'not detected'. †Denotes metabolites detected only in one replicate in MW at control conditions.

| | MW | BR | Y |
|---|---|---|---|
| Alanine | 1 ± 0,16 | 0,36 ± 0,06* | 2,37 ± 0,37* |
| Pyruvate | 1 ± 0,2 | 0,94 ± 0,13 | 0,76 ± 0,06 |
| Valine | 1 ± 0,19 | 0,64 ± 0,09* | 0,21 ± 0,03* |
| Glycerol | 1 ± 0,23 | 1,1 ± 0,18 | 1,35 ± 0,28 |
| Isoleucine | 1 ± 0,2 | 1,03 ± 0,13 | 0,67 ± 0,22 |
| Glycine | 1 ± 0,15 | 0,95 ± 0,19 | 0,78 ± 0,15 |
| Proline | 1 ± 0,16 | 0,59 ± 0,13 | 0,63 ± 0,38 |
| GABA | 1 ± 0,08 | 0,63 ± 0,13* | 0,42 ± 0,11* |
| Glycerate | 1 ± 0,09 | 1,07 ± 0,27 | 0,85 ± 0,1 |
| Serine | 1 ± 0,18 | 3,34 ± 0,95* | 1,55 ± 0,46 |
| Succinate | 1 ± 0,23 | 2,7 ± 0,55* | 1,83 ± 0,41 |
| Threonine | 1 ± 0,21 | 2,7 ± 0,51* | 0,93 ± 0,24 |
| Fumarate | 1 ± 0,12 | 0,88 ± 0,11 | 1,05 ± 0,2 |
| Nicotinate | 1 ± 0,14 | 1,13 ± 0,16 | 1,03 ± 0,18 |
| beta-Alanine | 1 ± 0,16 | 0,99 ± 0,13 | 0,14 ± 0,05* |
| Homoserine | 1 ± 0,13 | 2,18 ± 0,28* | 0,99 ± 0,22 |
| Methionine | 1 ± 0,23 | 3,83 ± 0,43* | 1,7 ± 0,49 |
| Glutamine | 1 ± 0,3 | 1,98 ± 0,46 | 0,78 ± 0,34 |
| Xylose | 1 ± 0,06 | 3,13 ± 0,68* | 4,38 ± 0,78* |
| Putrescine | 1 ± 0,16 | 2,3 ± 0,66 | 4,15 ± 0,94* |
| 2-oxo-Glutarate | 1 ± 0,1 | 2,49 ± 0,47* | 2,43 ± 0,34* |
| Phenylalanine | 1 ± 0,27 | 1,22 ± 0,39 | 0,77 ± 0,24 |
| Asparagine | 1 ± 0,32 | 3,77 ± 0,16* | 1,75 ± 0,54 |
| Ornithine | 1 ± 0,23 | 2,19 ± 0,32* | 0,74 ± 0,29 |
| Glycerol-3-P | 1† | 1,89 ± 0,24 | 2,07 ± 0,27 |
| Lysine | 1 ± 0,31 | 2,15 ± 0,35* | 0,57 ± 0,18 |
| Galacturonate | n.d. | 1 ± 0,28 | 12,33 ± 1,32* |
| Dehydroascorbate | 1 ± 0,07 | 1,34 ± 0,05* | 1,76 ± 0,14* |
| myo-Inositol | 1 ± 0,08 | 0,52 ± 0,08* | 0,55 ± 0,18* |
| Tyrosine | 1 ± 0,27 | 0,65 ± 0,24 | 0,12 ± 0,03* |
| Fructose-6-P | 1 ± 0,16 | 0,94 ± 0,15 | 1,47 ± 0,13* |
| Glucose-6-P | 1 ± 0,15 | 0,54 ± 0,08 | 0,8 ± 0,05 |
| Tryptophan | 1 ± 0,23 | 6,64 ± 2,38* | 1,39 ± 0,81 |
| Maltose | 1 ± 0,2 | 0,81 ± 0,19 | 1,4 ± 0,13 |
| Trehalose | n.d. | 1 ± 0,31 | 3,87 ± 0,74* |
| Galactinol | 1 ± 0,62 | n.d. | n.d. |
| 3-caffeoyl, cis-Quinate | 1 ± 0,15 | 0,84 ± 0,17 | 0,39† |
| 3-caffeoyl, trans-Quinate | 1 ± 0,13 | 1,77 ± 0,74 | 0,49† |
| Raffinose | 1 ± 0,06 | 3,64 ± 0,6* | 2,58 ± 0,51 |
| AMP | n.d. | 1 ± 0,2 | 4,05 ± 0,93* |
| Malate | 1 ± 0,07 | 0,6 ± 0,13* | 0,21 ± 0,06* |
| Aspartate | 1 ± 0,17 | 2,7 ± 0,58* | 4,32 ± 0,8* |
| Glutamate | 1 ± 0,13 | 8,74 ± 2,24* | 7,88 ± 1,71* |
| Fructose | 1 ± 0,07 | 0,9 ± 0,16 | 1,05 ± 0,21 |
| Citrate | 1 ± 0,14 | 2,36 ± 0,44* | 2,13 ± 0,28* |
| Glucose | 1 ± 0,12 | 0,85 ± 0,15 | 1,36 ± 0,28 |
| Sucrose | 1 ± 0,14 | 1,38 ± 0,29 | 1,08 ± 0,25 |

**Table S6 Relative metabolite levels from MG to MG+3 detected by GC-MS in tomato WT and *aox1a* mutant fruits.** Values were determined after normalization by the mean levels of WT pericarp samples at mature green (MG) stage. Levels correspond to the mean ± SE of n=4-6 independent replicates. Letters denote significant differences (P < 0.05) among genotypes at all ripening stages.

| Metabolite | WT_MG | WT_MG+1 | WT_MG+2 | WT_MG+3 | a1_MG | a1_MG+1 | a1_MG+2 | a1_MG+3 | a2_MG | a2_MG+1 | a2_MG+2 | a2_MG+3 |
|---|---|---|---|---|---|---|---|---|---|---|---|---|
| 2-amino-Adipic acid | 1 ± 0.17b | 0.62 ± 0.13ab | 0.63 ± 0.26ab | 0.38 ± 0.07a | 0.79 ± 0.22ab | 1 ± 0.19b | 0.44 ± 0.11a | 0.34 ± 0.07a | 1.03 ± 0.21b | 0.72 ± 0.13ab | 0.4 ± 0.08a | 0.47 ± 0.07a |
| 2-oxo-glutarate | 1 ± 0.16a | 2.09 ± 0.35ab | 1.3 ± 0.15ab | 1.98 ± 0.37ab | 1.34 ± 0.2ab | 1.33 ± 0.19ab | 2.96 ± 1.11b | 1.65 ± 0.3ab | 1.56 ± 0.45ab | 1.1 ± 0.17a | 1.61 ± 0.38ab | 2.55 ± 0.62ab |
| 3-deoxy-glucosone | 1 ± 0.14b | 1.08 ± 0.18b | 0.87 ± 0.2ab | 0.85 ± 0.08ab | 0.47 ± 0.14ab | 0.95 ± 0.11ab | 0.93 ± 0.12ab | 0.72 ± 0.07ab | 0.96 ± 0.21ab | 0.92 ± 0.13ab | 0.75 ± 0.23ab | 0.7 ± 0.09ab |
| Asparagine | 1 ± 0.15abc | 1.83 ± 0.48abc | 1.16 ± 0.26abc | 0.81 ± 0.16ab | 0.51 ± 0.14ab | 0.93 ± 0.12abc | 1.09 ± 0.07abc | 1.1 ± 0.18abc | 1.91 ± 0.7bc | 1.49 ± 0.45abc | 0.68 ± 0.21abc | 2.23 ± 0.89c |
| Aspartate | 1 ± 0.14a | 1.42 ± 0.25ab | 2.38 ± 1.02b | 1.18 ± 0.2a | 0.92 ± 0.15a | 1.21 ± 0.19a | 1.21 ± 0.16a | 1.07 ± 0.12a | 1.1 ± 0.2a | 1.05 ± 0.19a | 1.32 ± 0.09ab | 1.4 ± 0.31ab |
| Benzoate | 1 ± 0.15b | 0.92 ± 0.09ab | 0.77 ± 0.14ab | 0.97 ± 0.15b | 0.85 ± 0.06ab | 0.88 ± 0.14ab | 0.73 ± 0.15ab | 0.92 ± 0.12ab | 0.44 ± 0.06a | 1.01 ± 0.16b | 0.66 ± 0.13ab | 1.06 ± 0.35b |
| beta-Alanine | 1 ± 0.15a | 1.63 ± 0.35ab | 0.92 ± 0.21a | 0.84 ± 0.22a | 1.18 ± 0.36ab | 1.18 ± 0.11ab | 1.25 ± 0.22ab | 0.94 ± 0.11a | 1.82 ± 0.37b | 1.43 ± 0.23ab | 0.87 ± 0.25a | 0.94 ± 0.2a |
| Citrate | 1 ± 0.13abc | 1.86 ± 0.22cd | 1.5 ± 0.38bc | 1.5 ± 0.12bc | 0.55 ± 0.17a | 1.11 ± 0.1ab | 1.28 ± 0.17abc | 1.4 ± 0.19bc | 1.1 ± 0.22ab | 0.99 ± 0.14ab | 1.02 ± 0.4ab | 2.42 ± 0.2d |
| Dehydroascorbate | 1 ± 0.31a | 1.84 ± 0.65a | 1 ± 0.39a | 1.35 ± 0.22a | 0.67 ± 0.05a | 0.82 ± 0.09a | 1.65 ± 0.48a | 0.95 ± 0.17a | 0.66 ± 0.12a | 0.59 ± 0.13a | 0.84 ± 0.24a | 1.88 ± 0.72a |
| Erythritol | 1 ± 0.06ab | 1.27 ± 0.14abc | 1.21 ± 0.29abc | 1.71 ± 0.27cd | 0.67 ± 0.05a | 0.99 ± 0.08ab | 1.19 ± 0.15abc | 1.3 ± 0.16bc | 1.11 ± 0.19ab | 1.05 ± 0.12ab | 1.12 ± 0.28ab | 1.87 ± 0.15d |
| Fructose | 1 ± 0.09bcd | 1.17 ± 0.13cd | 0.87 ± 0.2ab | 1.34 ± 0.11d | 0.43 ± 0.12a | 0.89 ± 0.07bc | 1.03 ± 0.11bcd | 1.14 ± 0.11cd | 0.8 ± 0.15abc | 0.74 ± 0.1abc | 0.66 ± 0.18ab | 1.01 ± 0.17bcd |
| Fructose-6-P | 1 ± 0.32a | 0.84 ± 0.24a | 0.75 ± 0.29a | 0.9 ± 0.25a | 0.4 ± 0.2a | 0.84 ± 0.26a | 0.78 ± 0.19a | 0.78 ± 0.21a | 0.9 ± 0.26a | 0.68 ± 0.17a | 0.28 ± 0.08a | 0.79 ± 0.22a |
| Fucose | 1 ± 0.09bc | 1.01 ± 0.07bc | 0.9 ± 0.15abc | 1.21 ± 0.14c | 0.54 ± 0.13a | 0.94 ± 0.02bc | 0.96 ± 0.07bc | 1.01 ± 0.1bc | 0.87 ± 0.15abc | 0.86 ± 0.07abc | 0.74 ± 0.19ab | 1.02 ± 0.06bc |
| Fumarate | 1 ± 0.12b | 0.77 ± 0.1ab | 0.69 ± 0.16ab | 1.06 ± 0.2b | 0.38 ± 0.12a | 0.63 ± 0.05ab | 0.7 ± 0.08ab | 0.77 ± 0.1ab | 0.83 ± 0.19ab | 0.75 ± 0.06ab | 0.64 ± 0.19ab | 0.89 ± 0.12b |
| GABA | 1 ± 0.09ab | 1.07 ± 0.07b | 0.86 ± 0.15ab | 0.85 ± 0.12ab | 0.73 ± 0.16ab | 1 ± 0.03ab | 0.94 ± 0.06ab | 0.9 ± 0.05ab | 0.96 ± 0.12ab | 0.93 ± 0.07ab | 0.69 ± 0.16a | 0.74 ± 0.08ab |
| Galactonic acid-1,4-lactone | 1 ± 0.18ab | 1.29 ± 0.21ab | 1.75 ± 0.68b | 1.75 ± 0.47b | 0.5 ± 0.13a | 1.05 ± 0.15ab | 0.88 ± 0.17ab | 0.97 ± 0.2ab | 0.96 ± 0.07ab | 0.74 ± 0.12ab | 0.8 ± 0.19ab | 1.58 ± 0.42b |
| Glucarate | 1 ± 0.34a | 0.94 ± 0.27a | 0.32 ± 0.1a | 0.66 ± 0.2a | 0.24 ± 0.1a | 0.83 ± 0.29a | 0.72 ± 0.19a | 0.52 ± 0.12a | 0.75 ± 0.15a | 0.66 ± 0.21a | 0.39 ± 0.26a | 0.46 ± 0.08a |
| Glucose | 1 ± 0.09ab | 1.14 ± 0.13bc | 0.91 ± 0.2ab | 1.53 ± 0.25c | 0.58 ± 0.11a | 0.87 ± 0.05ab | 1.01 ± 0.12ab | 1.09 ± 0.09b | 0.78 ± 0.14ab | 0.75 ± 0.07ab | 0.66 ± 0.18ab | 0.93 ± 0.2ab |
| Glucose-6-P | 1 ± 0.34a | 1.07 ± 0.39a | 0.65 ± 0.31a | 1.11 ± 0.36a | 0.57 ± 0.21a | 0.95 ± 0.31a | 0.92 ± 0.36a | 0.91 ± 0.31a | 1.51 ± 0.6a | 0.8 ± 0.3a | 0.45 ± 0.16a | 0.92 ± 0.31a |
| Glutamate | 1 ± 0.18a | 1.27 ± 0.11a | 0.91 ± 0.17a | 1.19 ± 0.15a | 0.88 ± 0.15a | 1.29 ± 0.13a | 1.59 ± 0.27ab | 1.35 ± 0.14a | 1.4 ± 0.27a | 1.21 ± 0.21a | 1.44 ± 0.12ab | 2.05 ± 0.38b |
| Glutamine | 1 ± 0.26a | 1.35 ± 0.33a | 1.1 ± 0.35a | 0.87 ± 0.24a | 0.93 ± 0.28a | 0.87 ± 0.18a | 1.49 ± 0.54a | 1.11 ± 0.22a | 1.55 ± 0.8a | 1.35 ± 0.14a | 1.07 ± 0.35a | 1.78 ± 0.41a |
| Glycerate | 1 ± 0.12c | 0.72 ± 0.08abc | 0.87 ± 0.21bc | 1.62 ± 0.16d | 0.41 ± 0.08a | 0.76 ± 0.09abc | 0.67 ± 0.04abc | 0.87 ± 0.05bc | 0.87 ± 0.15bc | 0.95 ± 0.11bc | 0.58 ± 0.07ab | 1.08 ± 0.16c |
| Glycerol | 1 ± 0.29ab | 0.69 ± 0.08ba | 1.29 ± 0.56ab | 0.74 ± 0.15ab | 0.6 ± 0.15a | 0.81 ± 0.13ab | 1.01 ± 0.1ab | 0.73 ± 0.09ab | 1.25 ± 0.64ab | 0.81 ± 0.11ab | 0.92 ± 0.33ab | 1.72 ± 0.49b |
| Glycerol-3-P | 1 ± 0.1a | 1.16 ± 0.13a | 0.93 ± 0.14a | 1 ± 0.15a | 0.94 ± 0.11a | 1.24 ± 0.12a | 1.24 ± 0.09a | 1.04 ± 0.08a | 1.12 ± 0.11a | 1.14 ± 0.04a | 0.96 ± 0.15a | 0.94 ± 0.08a |
| Glycine | 1 ± 0.16ab | 1.22 ± 0.13ab | 0.95 ± 0.2ab | 0.94 ± 0.09ab | 1.58 ± 0.37ab | 1.2 ± 0.14ab | 1.3 ± 0.23ab | 1.41 ± 0.26b | 1.33 ± 0.23a | 1.07 ± 0.13ab | 0.81 ± 0.23ab | 1.05 ± 0.24ab |
| Histidine | 1 ± 0.2a | 2.82 ± 0.57c | 2.35 ± 0.48abc | 1.55 ± 0.36abc | 0.75 ± 0.2a | 0.89 ± 0.11a | 1.5 ± 0.24abc | 1.67 ± 0.23abc | 2.03 ± 1.02abc | 1.11 ± 0.3ab | 1.15 ± 0.2abc | 2.73 ± 0.73bc |
| Homoserine | 1 ± 0.19a | 1.22 ± 0.25a | 1.09 ± 0.31a | 0.67 ± 0.09a | 0.84 ± 0.15a | 1.12 ± 0.32a | 0.7 ± 0.18a | 0.66 ± 0.06a | 0.77 ± 0.1a | 1.09 ± 0.14a | 1.01 ± 0.23a | 0.91 ± 0.08a |
| Isocitrate | 1 ± 0.09a | 1.34 ± 0.07a | 1.76 ± 0.16a | 1.33 ± 0.11a | 1.07 ± 0.18a | 1.46 ± 0.33a | 1.33 ± 0.1a | 1.43 ± 0.09a | 1.39 ± 0.01a | 1 ± 0.04a | 1.45 ± 0.27a | 1.6 ± 0.07a |
| Isoleucine | 1 ± 0.12a | 1.92 ± 0.52b | 0.94 ± 0.19a | 1.06 ± 0.21ab | 0.77 ± 0.23a | 0.91 ± 0.05a | 1.51 ± 0.5ab | 0.89 ± 0.02a | 1.58 ± 0.35ab | 1.17 ± 0.22ab | 0.89 ± 0.25a | 1.15 ± 0.18ab |
| Isomaltose | 1 ± 0.14a | 1.47 ± 0.25a | 0.54 ± 0.11a | 1.2 ± 0.09a | 0.74 ± 0.21a | 1.21 ± 0.23a | 1.53 ± 0.38a | 1.17 ± 0.14a | 1.35 ± 0.34a | 0.95 ± 0.14a | 1.16 ± 0.4a | 1.08 ± 0.06a |
| Lysine | 1 ± 0.05ab | 0.92 ± 0.04ab | 1.57 ± 0.23bc | 1.12 ± 0.12abc | 0.58 ± 0.05a | 1.11 ± 0.18abc | 1.46 ± 0.16bc | 1.06 ± 0.16ab | 1.26 ± 0.31abc | 1.02 ± 0.2ab | 0.94 ± 0ab | 1.82 ± 0.17c |
| Malate | 1 ± 0.13bc | 1.24 ± 0.16c | 1.12 ± 0.24bc | 1.19 ± 0.19bc | 0.36 ± 0.1a | 0.72 ± 0.05ab | 0.79 ± 0.11abc | 0.82 ± 0.14abc | 0.86 ± 0.17abc | 0.79 ± 0.13abc | 0.72 ± 0.2ab | 1.74 ± 0.16d |
| Maltose | 1 ± 0.16ab | 0.99 ± 0.07ab | 1.3 ± 0.28ab | 1.79 ± 0.1b | 0.62 ± 0.09a | 1.21 ± 0.27ab | 1.35 ± 0.29ab | 1.26 ± 0.14ab | 1.57 ± 0.34ab | 0.96 ± 0.25ab | 0.95 ± 0.23ab | 1.25 ± 0.36ab |
| Methionine | 1 ± 0.05ab | 2.09 ± 0.32bc | 1.6 ± 0.41abc | 1.06 ± 0.16ab | 1.4 ± 0.26abc | 0.73 ± 0.06a | 1.56 ± 0.31abc | 0.97 ± 0.09ab | 1.69 ± 0.46abc | 1.15 ± 0.01ab | 1.36 ± 0.14abc | 2.49 ± 0.56c |
| myo-Inositol | 1 ± 0.07a | 1.03 ± 0.06a | 0.81 ± 0.16a | 1.16 ± 0.13a | 0.77 ± 0.19a | 1.02 ± 0.07a | 0.99 ± 0.08a | 0.97 ± 0.09a | 1.05 ± 0.19a | 0.97 ± 0.08a | 1.09 ± 0.07a | 0.94 ± 0.1a |
| Nicotinate | 1 ± 0.08ab | 1.24 ± 0.07bc | 1.23 ± 0.22bc | 1.58 ± 0.19c | 0.59 ± 0.12a | 1.01 ± 0.1ab | 1.12 ± 0.07bc | 1.15 ± 0.11bc | 0.95 ± 0.17ab | 0.99 ± 0.06ab | 1.12 ± 0.2bc | 1.42 ± 0.18bc |
| Ornithine | 1 ± 0.1a | 1.08 ± 0.09ab | 1.04 ± 0.11ab | 0.94 ± 0.1ab | 1.63 ± 0.62a | 1.06 ± 0.08a | 1.08 ± 0.11ab | 1.18 ± 0.19ab | 1.33 ± 0.18ab | 1 ± 0.15ab | 1.02 ± 0.17ab | 1.66 ± 0.19b |
| Phenylalanine | 1 ± 0.24ab | 1.89 ± 0.4b | 0.83 ± 0.22ab | 0.65 ± 0.08a | 0.91 ± 0.28ab | 0.95 ± 0.05ab | 1.49 ± 0.64ab | 0.99 ± 0.25ab | 1.26 ± 0.31ab | 0.96 ± 0.26ab | 0.95 ± 0.36ab | 1 ± 0.18ab |
| Phosphoric acid | 1 ± 0.08b | 1.06 ± 0.07b | 0.88 ± 0.12b | 0.99 ± 0.04b | 0.6 ± 0.13a | 0.93 ± 0.04b | 0.91 ± 0.04b | 0.92 ± 0.04b | 0.92 ± 0.11b | 0.84 ± 0.06ab | 0.84 ± 0.13ab | 0.97 ± 0.03b |
| Proline | 1 ± 0.17a | 2.25 ± 0.53ab | 1.12 ± 0.28ab | 2.57 ± 0.79ab | 4.14 ± 1.14ab | 5.48 ± 0.76bc | 8.7 ± 3.83c | 3.42 ± 0.69ab | 2.08 ± 0.48ab | 2.05 ± 0.36ab | 2.41 ± 0.66ab | 2.39 ± 0.45ab |
| Putrescine | 1 ± 0.09a | 1.36 ± 0.09a | 1.03 ± 0.25a | 1.29 ± 0.26a | 0.69 ± 0.1a | 1 ± 0.12a | 1.11 ± 0.13a | 1.37 ± 0.26a | 1.12 ± 0.3a | 1.11 ± 0.19a | 0.71 ± 0.2a | 2.54 ± 0.27b |
| Pyruvate | 1 ± 0.13abc | 1.56 ± 0.24abcd | 0.98 ± 0.23ab | 1.63 ± 0.43abcd | 0.91 ± 0.24ab | 1.13 ± 0.16abcd | 2.23 ± 0.59d | 1.46 ± 0.2abcd | 0.68 ± 0.08a | 0.72 ± 0.15a | 2.13 ± 0.62cd | 1.94 ± 0.28bcd |
| Quinate | 1 ± 0.07c | 0.34 ± 0.02ab | 0.45 ± 0.12ab | 0.49 ± 0.11ab | 0.79 ± 0.01bc | 0.64 ± 0.16abc | 0.36 ± 0.09ab | 0.23 ± 0.04a | 0.37 ± 0.06ab | 0.55 ± 0abc | 0.24 ± 0.03a | 0.45 ± 0.07ab |
| Rhamnose | 1 ± 0.07b | 1.15 ± 0.11bc | 1.02 ± 0.23b | 1.31 ± 0.17bc | 0.51 ± 0.11a | 0.93 ± 0.03ab | 1.08 ± 0.14bc | 1.06 ± 0.12bc | 1 ± 0.2b | 0.85 ± 0.07ab | 0.86 ± 0.26ab | 1.53 ± 0.09c |
| Serine | 1 ± 0.2ab | 2.12 ± 0.45c | 1.06 ± 0.21ab | 1.84 ± 0.31bc | 0.7 ± 0.13a | 1.83 ± 0.31bc | 2.28 ± 0.24c | 3.22 ± 0.28d | 1.61 ± 0.33abc | 1.13 ± 0.15ab | 1.08 ± 0.33ab | 2.2 ± 0.37c |
| Spermidine | 1 ± 0.13ab | 1.24 ± 0.16abc | 1.61 ± 0.15bc | 1.04 ± 0.17ab | 0.66 ± 0.1a | 0.71 ± 0.09a | 1.08 ± 0.06abc | 0.93 ± 0.03a | 1.62 ± 0.15bc | 1.01 ± 0.02ab | 1.31 ± 0.29abc | 1.71 ± 0.13c |
| Succinate | 1 ± 0.11a | 0.7 ± 0.13a | 0.78 ± 0.09a | 0.75 ± 0.14a | 0.76 ± 0.18a | 0.75 ± 0.05a | 0.53 ± 0.04a | 0.57 ± 0.05a | 0.72 ± 0.21a | 1.02 ± 0.16a | 0.55 ± 0.02a | 0.92 ± 0.09a |
| Sucrose | 1 ± 0.08ab | 1.01 ± 0.09ab | 0.67 ± 0.16a | 1.57 ± 0.29bcd | 0.64 ± 0.17a | 1.77 ± 0.2cd | 2.16 ± 0.49d | 1.93 ± 0.1cd | 0.91 ± 0.16ab | 1.28 ± 0.13abc | 0.97 ± 0.26ab | 1.36 ± 0.15abc |
| Threonine | 1 ± 0.14ab | 2.16 ± 0.66c | 0.95 ± 0.23a | 0.85 ± 0.1a | 0.53 ± 0.14a | 0.74 ± 0.04a | 1.02 ± 0.18ab | 0.75 ± 0.05a | 1.97 ± 0.57bc | 1.34 ± 0.29abc | 1.02 ± 0.29ab | 1.1 ± 0.19ab |
| Trehalose | 1 ± 0.13abc | 1.52 ± 0.21bc | 0.59 ± 0.14a | 1.78 ± 0.33c | 0.68 ± 0.02ab | 1.24 ± 0.15abc | 1.62 ± 0.37c | 1.52 ± 0.22bc | 1.63 ± 0.13c | 1.03 ± 0.17abc | 1.58 ± 0.27bc | 1.33 ± 0.09abc |
| Tryptophan | 1 ± 0.24abc | 4.54 ± 0.42c | 0.51 ± 0.17a | 1.92 ± 0.45bc | 0.32 ± 0.09a | 0.69 ± 0.08a | 1.38 ± 0.4abc | 0.85 ± 0.17ab | 1.23 ± 0.39abc | 0.75 ± 0.17a | 0.48 ± 0.18a | 2.1 ± 0.18c |
| Tyramine | 1 ± 0.14ab | 1.28 ± 0.24abcc | 0.74 ± 0.19a | 1.63 ± 0.17bcde | 0.71 ± 0.17a | 2.29 ± 0.18e | 1.9 ± 0.3de | 1.83 ± 0.31cde | 1.25 ± 0.28abcd | 1.09 ± 0.06abc | 1.11 ± 0.29abc | 1.86 ± 0.19de |
| Tyrosine | 1 ± 0.12ab | 1.09 ± 0.21ab | 1.16 ± 0.25ab | 1.02 ± 0.29ab | 0.42 ± 0.14a | 0.95 ± 0.06ab | 1.4 ± 0.4ab | 0.73 ± 0.04b | 1.26 ± 0.23ab | 0.99 ± 0.26ab | 0.41 ± 0.14a | 0.94 ± 0.19ab |
| Urea | 1 ± 0.15b | 0.81 ± 0.05ab | 0.85 ± 0.18ab | 0.71 ± 0.09ab | 0.53 ± 0.08a | 0.81 ± 0.16ab | 0.86 ± 0.09ab | 0.72 ± 0.06ab | 0.7 ± 0.09ab | 0.59 ± 0.01ab | 0.6 ± 0.08ab | 0.71 ± 0.06ab |
| Valine | 1 ± 0.16a | 1.37 ± 0.24a | 0.84 ± 0.19a | 0.94 ± 0.2a | 0.85 ± 0.25a | 1.01 ± 0.11a | 1.27 ± 0.22a | 1.05 ± 0.07a | 1.26 ± 0.24a | 1.05 ± 0.17a | 0.95 ± 0.25a | 1.05 ± 0.37a |
| Xylose | 1 ± 0.1ab | 1.09 ± 0.09abc | 1.2 ± 0.42abc | 1.9 ± 0.18c | 0.52 ± 0.1a | 0.84 ± 0.03ab | 1.24 ± 0.17abc | 1.32 ± 0.15abc | 0.88 ± 0.21ab | 0.8 ± 0.11ab | 1.13 ± 0.28abc | 1.58 ± 0.35bc |

**Table S7 Relative metabolite levels from MG+5 to MG+13 stages detected by GC-MS in tomato WT and aox1a mutant fruits.** Values were determined after normalization by the mean levels of WT pericarp samples at mature green (MG) stage. Levels correspond to the mean ± SE of n=4-6 independent replicates. Letters denote significant differences (P < 0.05) among genotypes at all ripening stages.

| Metabolite | WT_MG+5 | WT_MG+7 | WT_MG+9 | WT_MG+11 | WT_MG+13 | a1_MG+5 | a1_MG+7 | a1_MG+9 | a1_MG+11 | a1_MG+13 | a2_MG+5 | a2_MG+7 | a2_MG+9 | a2_MG+11 | a2_MG+13 |
|---|---|---|---|---|---|---|---|---|---|---|---|---|---|---|---|
| 2-amino-Adipic acid | 0.62 ± 0.1ab | 0.82 ± 0.14b | 0.64 ± 0.07ab | 0.69 ± 0.09ab | 0.48 ± 0.1ab | 0.37 ± 0.04a | 0.58 ± 0.12ab | 0.63 ± 0.11ab | 0.78 ± 0.18b | 0.8 ± 0.12b | 0.77 ± 0.06b | 0.49 ± 0.09ab | 0.56 ± 0.13ab | 0.6 ± 0.11ab | 0.57 ± 0.04ab |
| 2-oxo-glutarate | 4.3 ± 1.1abc | 5.52 ± 1.02abc | 4.9 ± 0.18abc | 5.81 ± 0.62bc | 4.32 ± 0.37abc | 2.73 ± 0.53a | 2.71 ± 0.4a | 3.24 ± 0.45ab | 8.89 ± 0.68d | 6.76 ± 0.73cd | 4.75 ± 0.87abc | 3.83 ± 0.74ab | 4.26 ± 0.79ab | 4.97 ± 1.17abc | 8.6 ± 1.52d |
| 3-deoxy-glucosone | 0.75 ± 0.17ab | 1.13 ± 0.12ab | 0.79 ± 0.12a | 1.14 ± 0.19ab | 1.16 ± 0.11ab | 0.79 ± 0.12a | 0.78 ± 0.1a | 0.9 ± 0.2a | 1.04 ± 0.19ab | 1.47 ± 0.15b | 0.7 ± 0.13a | 0.88 ± 0.16a | 0.82 ± 0.2a | 0.93 ± 0.13a | 1.52 ± 0.26b |
| Asparagine | 0.85 ± 0.23a | 1.93 ± 0.59ab | 0.55 ± 0.24a | 0.67 ± 0.12a | 1.14 ± 0.33a | 1.44 ± 0.39a | 2.08 ± 0.78ab | 2.37 ± 0.74ab | 0.96 ± 0.2a | 2.1 ± 0.78ab | 1.87 ± 0.6ab | 0.96 ± 0.42a | 2.1 ± 0.82ab | 1.86 ± 0.36ab | 3.53 ± 0.94b |
| Aspartate | 13.11 ± 3.83bc | 3.92 ± 1.31a | 9.44 ± 1.91bc | 19.36 ± 6.39b | 30.29 ± 9.33c | 2.04 ± 0.57a | 1.86 ± 0.35a | 2.59 ± 0.71a | 3.67 ± 0.65a | 5.15 ± 1.18a | 1.83 ± 0.63a | 1.21 ± 0.15a | 1.88 ± 0.17a | 1.76 ± 0.44a | 7.32 ± 1.2a |
| Benzoate | 1.09 ± 0.16a | 0.75 ± 0.12a | 1.08 ± 0.15a | 1.33 ± 0.3a | 1.31 ± 0.42a | 1.39 ± 0.39a | 0.76 ± 0.11a | 0.94 ± 0.23a | 1.17 ± 0.23a | 1.16 ± 0.2a | 1.1 ± 0.12a | 0.81 ± 0.12a | 1.11 ± 0.29a | 0.91 ± 0.07a | 1.4 ± 0.49a |
| beta-Alanine | 0.6 ± 0.23abc | 0.62 ± 0.17abc | 0.22 ± 0.02a | 0.23 ± 0.03a | 0.24 ± 0.07a | 0.97 ± 0.16bc | 1.06 ± 0.22cd | 1.57 ± 0.47d | 0.3 ± 0.02a | 0.32 ± 0.04a | 0.74 ± 0.16abc | 0.77 ± 0.23abc | 0.83 ± 0.27abc | 0.53 ± 0.11abc | 0.4 ± 0.09ab |
| Citrate | 2.74 ± 0.45abcd | 2.55 ± 0.39abcd | 3.3 ± 0.15bcd | 3.6 ± 0.09d | 3.34 ± 0.2cd | 2.24 ± 0.33ab | 2.39 ± 0.4abc | 2.53 ± 0.36abcd | 3.27 ± 0.03bcd | 3.29 ± 0.36bcd | 2.73 ± 0.22abcd | 2.02 ± 0.31a | 2.75 ± 0.39abcd | 2.96 ± 0.51abcd | 3.54 ± 0.27d |
| Dehydroascorbate | 8.32 ± 5.1abc | 11.84 ± 2.53abcd | 16.12 ± 5.14abcd | 23.32 ± 2.79d | 11.48 ± 5.49abcd | 3.97 ± 1.04ab | 3.12 ± 1.65a | 12.98 ± 1.2abcd | 9.43 ± 3.42abcd | 18.66 ± 2cd | 8.37 ± 2.56abc | 14.02 ± 3.77abcd | 8.47 ± 1.14abc | 19.99 ± 2.91cd | 18.26 ± 2.1bcd |
| Erythritol | 1.52 ± 0.25ab | 1.77 ± 0.13abcd | 2.08 ± 0.13bcd | 2.31 ± 0.12cde | 2.02 ± 0.17abcd | 1.89 ± 0.3abcd | 1.42 ± 0.22a | 1.54 ± 0.22ab | 2.39 ± 0.24de | 2.37 ± 0.23de | 2.15 ± 0.22bcd | 1.39 ± 0.12a | 1.67 ± 0.15abcb | 1.96 ± 0.13abcd | 2.85 ± 0.19e |
| Fructose | 1.22 ± 0.19ab | 1.45 ± 0.07ab | 1.5 ± 0.09b | 1.49 ± 0.07b | 1.27 ± 0.06ab | 1.23 ± 0.09ab | 1.04 ± 0.17a | 1.18 ± 0.14ab | 1.48 ± 0.07b | 1.47 ± 0.1b | 1.35 ± 0.11ab | 1.32 ± 0.13ab | 1.3 ± 0.18ab | 1.36 ± 0.1ab | 1.3 ± 0.2ab |
| Fructose-6-P | 0.57 ± 0.15a | 0.81 ± 0.24a | 0.94 ± 0.38a | 0.84 ± 0.23a | 0.71 ± 0.18a | 1.03 ± 0.31a | 0.68 ± 0.2a | 0.84 ± 0.27a | 0.9 ± 0.16a | 1 ± 0.22a | 0.82 ± 0.21a | 0.74 ± 0.25a | 0.65 ± 0.19a | 0.71 ± 0.2a | 1.21 ± 0.29a |
| Fucose | 1.06 ± 0.14ab | 1.16 ± 0.08abc | 1.34 ± 0.04abcd | 1.48 ± 0.05cd | 1.32 ± 0.14abcd | 1.24 ± 0.13abc | 1 ± 0.14a | 1.16 ± 0.06abc | 1.44 ± 0.06bcd | 1.66 ± 0.18de | 1.24 ± 0.08abc | 1.07 ± 0.04ab | 1.18 ± 0.07abc | 1.39 ± 0.09abcd | 1.88 ± 0.29e |
| Fumarate | 0.78 ± 0.17abc | 0.94 ± 0.11bc | 0.46 ± 0.08a | 0.67 ± 0.05abc | 0.7 ± 0.08abc | 0.96 ± 0.21bc | 0.82 ± 0.18abc | 0.99 ± 0.17c | 0.53 ± 0.04ab | 0.54 ± 0.09abc | 0.91 ± 0.18bc | 0.93 ± 0.09bc | 0.81 ± 0.11abc | 0.89 ± 0.18abc | 0.72 ± 0.06abc |
| GABA | 0.62 ± 0.16abc | 0.8 ± 0.1bc | 0.34 ± 0.06a | 0.41 ± 0.07a | 0.4 ± 0.12a | 0.96 ± 0.09c | 0.85 ± 0.06bc | 0.93 ± 0.12bc | 0.57 ± 0.12ab | 0.61 ± 0.15abc | 0.71 ± 0.09abc | 0.61 ± 0.11abc | 0.68 ± 0.15abc | 0.7 ± 0.13abc | 0.78 ± 0.08bc |
| Galactonic acid-1,4-lactone | 3.45 ± 1.34abc | 3.17 ± 0.95abc | 5.57 ± 1.52abc | 5.51 ± 1.25abc | 4.39 ± 1.38abc | 2.32 ± 1.04abc | 2.82 ± 1.12abc | 1.75 ± 0.27a | 4.3 ± 1.38abcd | 6.09 ± 1.91c | 3.58 ± 0.77abc | 2.02 ± 0.42ab | 2.48 ± 0.48abc | 2.9 ± 0.37abc | 5.78 ± 1.55bc |
| Glucarate | 0.57 ± 0.15a | 0.66 ± 0.1a | 0.81 ± 0.17a | 0.76 ± 0.11a | 0.45 ± 0.08a | 0.8 ± 0.21a | 0.47 ± 0.1a | 0.59 ± 0.09a | 0.76 ± 0.1a | 0.74 ± 0.13a | 0.86 ± 0.13a | 0.9 ± 0.24a | 0.68 ± 0.24a | 0.7 ± 0.1a | 0.87 ± 0.24a |
| Glucose | 1.17 ± 0.21a | 1.42 ± 0.16a | 1.41 ± 0.05a | 1.22 ± 0.09a | 0.95 ± 0.1a | 1.14 ± 0.07a | 0.96 ± 0.19a | 0.98 ± 0.12a | 1.46 ± 0.25a | 1.28 ± 0.17a | 1.23 ± 0.11a | 1.38 ± 0.17a | 1.17 ± 0.23a | 1.18 ± 0.11a | 1.22 ± 0.37a |
| Glucose-6-P | 0.89 ± 0.3a | 1 ± 0.33a | 0.72 ± 0.22a | 1.02 ± 0.4a | 0.86 ± 0.29a | 1.01 ± 0.33a | 0.57 ± 0.24a | 0.95 ± 0.37a | 1.3 ± 0.45a | 0.95 ± 0.21a | 1.21 ± 0.42a | 0.68 ± 0.2a | 0.83 ± 0.29a | 1.07 ± 0.4a | 1.51 ± 0.54a |
| Glutamate | 1.32 ± 0.2a | 3.53 ± 1.49a | 15.51 ± 9.12a | 11.36 ± 6.55a | 2.46 ± 0.51a | 9.12 ± 6.75a | 3.61 ± 2.08a | 5.34 ± 2.56a | 28.82 ± 7ab | 52.6 ± 13.24b | 2.86 ± 1.6a | 1.48 ± 0.37a | 3.49 ± 0.97a | 5.79 ± 2.88a | 50.75 ± 21.55b |
| Glutamine | 0.87 ± 0.2a | 3.21 ± 1.5b | 1.03 ± 0.22a | 0.83 ± 0.2a | 0.97 ± 0.21a | 1.28 ± 0.28ab | 2.88 ± 0.73ab | 2.3 ± 0.72ab | 1.21 ± 0.37ab | 1.47 ± 0.35ab | 2.35 ± 0.9ab | 1.17 ± 0.33ab | 1.95 ± 0.43ab | 1.87 ± 0.37ab | 2.19 ± 0.43ab |
| Glycerate | 0.56 ± 0.11a | 0.71 ± 0.1ab | 0.53 ± 0.05a | 0.51 ± 0.04a | 0.59 ± 0.08ab | 0.8 ± 0.11ab | 0.62 ± 0.06ab | 0.69 ± 0.1ab | 0.63 ± 0.07ab | 0.57 ± 0.04a | 0.74 ± 0.06ab | 0.89 ± 0.16bc | 0.54 ± 0.05a | 1.11 ± 0.1c | 0.69 ± 0.08ab |
| Glycerol | 0.57 ± 0.14a | 0.58 ± 0.06a | 0.92 ± 0.19ab | 1.27 ± 0.26ab | 1.15 ± 0.45ab | 0.94 ± 0.38ab | 0.8 ± 0.32a | 0.8 ± 0.28a | 2.48 ± 0.63c | 1.22 ± 0.15ab | 2.05 ± 0.83bc | 0.51 ± 0.1a | 0.69 ± 0.25a | 0.71 ± 0.15a | 1.17 ± 0.17ab |
| Glycerol-3-P | 0.73 ± 0.08a | 0.98 ± 0.1abc | 0.86 ± 0.11a | 0.9 ± 0.08ab | 0.75 ± 0.06a | 1.25 ± 0.12cd | 1.01 ± 0.07abc | 1.2 ± 0.08bcd | 0.99 ± 0.09abc | 1.02 ± 0.08abc | 0.87 ± 0.07ab | 0.99 ± 0.12abc | 0.97 ± 0.09abc | 1.04 ± 0.07abc | 1.43 ± 0.2d |
| Glycine | 0.69 ± 0.14a | 0.82 ± 0.18a | 0.73 ± 0.15a | 0.65 ± 0.04a | 0.72 ± 0.11a | 1.18 ± 0.15a | 0.86 ± 0.22a | 1.17 ± 0.24a | 1.05 ± 0.17a | 1.27 ± 0.37ab | 0.76 ± 0.16a | 0.71 ± 0.21a | 0.74 ± 0.13a | 0.79 ± 0.19a | 1.82 ± 0.34b |
| Histidine | 2.79 ± 0.86a | 7.42 ± 3.45a | 5.14 ± 1.99a | 6.46 ± 1.44a | 9.16 ± 2.65a | 2.99 ± 0.88a | 2.72 ± 0.5a | 3.66 ± 0.65a | 4.52 ± 0.61a | 21.12 ± 4.61b | 5.86 ± 1.05a | 1.36 ± 0.33a | 3.53 ± 0.57a | 9.51 ± 2.64a | 30.09 ± 6.2c |
| Homoserine | 0.93 ± 0.09a | 1.04 ± 0.12ab | 1.19 ± 0.18ab | 1.24 ± 0.22ab | 1.03 ± 0.14ab | 1.04 ± 0.22ab | 1.05 ± 0.06ab | 1.07 ± 0.08ab | 1.11 ± 0.07ab | 1.2 ± 0.2ab | 1.01 ± 0.11ab | 0.91 ± 0.09a | 1 ± 0.08a | 1.1 ± 0.14ab | 1.51 ± 0.24b |
| Isocitrate | 2.03 ± 0.08abcd | 1.25 ± 0.14a | 2.1 ± 0.01bcd | 2.26 ± 0.04cd | 2.29 ± 0.01cd | 1.87 ± 0.28abc | 1.79 ± 0.23abc | 1.48 ± 0.24abc | 2.11 ± 0.06bcd | 2.84 ± 0.17d | 1.66 ± 0.02abc | 1.26 ± 0.21a | 1.56 ± 0.07abc | 1.28 ± 0.01ab | 2.74 ± 0.07d |
| Isoleucine | 0.54 ± 0.13a | 0.83 ± 0.31ab | 0.48 ± 0.1a | 0.51 ± 0.06a | 0.62 ± 0.13a | 1.03 ± 0.18ab | 0.98 ± 0.28ab | 0.97 ± 0.24ab | 0.6 ± 0.11a | 0.98 ± 0.35ab | 0.87 ± 0.15ab | 0.61 ± 0.15a | 0.78 ± 0.2ab | 0.76 ± 0.1ab | 1.31 ± 0.19b |
| Isomaltose | 1.49 ± 0.15a | 1.84 ± 0.25ab | 2.3 ± 0.32abc | 2.78 ± 0.26abc | 2.93 ± 0.31abc | 3.62 ± 1.12bc | 1.32 ± 0.35a | 1.38 ± 0.16a | 3.81 ± 0.63c | 3.53 ± 0.95bc | 1.79 ± 0.26ab | 1.23 ± 0.19a | 1.3 ± 0.07a | 1.67 ± 0.1ab | 4.2 ± 0.38c |
| Lysine | 1.53 ± 0.23ab | 1.91 ± 0.52ab | 5.79 ± 1.93bc | 5.07 ± 1.22abc | 4.1 ± 1.24ab | 1.71 ± 0.13ab | 2.82 ± 0.42ab | 2.83 ± 0.59ab | 2.86 ± 0.67ab | 9.24 ± 2.08c | 3.54 ± 0.52ab | 1.07 ± 0.18a | 2.5 ± 0.32ab | 5.78 ± 1.19bc | 15.76 ± 1.96d |
| Malate | 1.29 ± 0.4cd | 1.07 ± 0.21abcd | 0.6 ± 0.18abc | 0.49 ± 0.05ab | 0.76 ± 0.26abc | 1.05 ± 0.18abcd | 1.21 ± 0.22cd | 1.29 ± 0.2cd | 0.61 ± 0.14abc | 0.41 ± 0.07a | 1.51 ± 0.19d | 1.16 ± 0.12bcd | 1.18 ± 0.17bcd | 1.09 ± 0.25abcd | 0.59 ± 0.26abc |
| Maltose | 1.7 ± 0.23a | 1.88 ± 0.34a | 2.01 ± 0.31a | 2.18 ± 0.4ab | 2.2 ± 0.46ab | 2.27 ± 0.17ab | 1.24 ± 0.38a | 1.93 ± 0.62a | 3.04 ± 0.67ab | 2.2 ± 0.23ab | 2.33 ± 0.43ab | 1.33 ± 0.23a | 1.39 ± 0.26a | 1.95 ± 0.42a | 4.22 ± 1.48b |
| Methionine | 3.56 ± 0.74ab | 6.61 ± 2.36b | 5.72 ± 0.76b | 7.18 ± 0.85b | 6.24 ± 1.42b | 3.44 ± 1.33ab | 3.15 ± 0.71ab | 3.77 ± 0.59ab | 5.37 ± 0.55ab | 6.5 ± 0.97b | 5.62 ± 1.46b | 1.3 ± 0.09a | 4.1 ± 0.64ab | 5.51 ± 1.72ab | 12.62 ± 1.74c |
| myo-Inositol | 0.91 ± 0.1abc | 1.06 ± 0.14abcde | 0.98 ± 0.1abcd | 0.96 ± 0.08abcd | 0.83 ± 0.07a | 1.1 ± 0.06abcde | 0.9 ± 0.11ab | 1.19 ± 0.12bcde | 1.15 ± 0.06bcde | 1.24 ± 0.11de | 1.01 ± 0.1abcde | 1.17 ± 0.1bcde | 1.05 ± 0.07abcde | 1.23 ± 0.06cde | 1.31 ± 0.12e |
| Nicotinate | 1.51 ± 0.25abc | 1.54 ± 0.03abc | 1.77 ± 0.12abc | 2.09 ± 0.11c | 1.67 ± 0.16abc | 1.7 ± 0.16abc | 1.28 ± 0.16a | 1.64 ± 0.22abc | 1.91 ± 0.18bcd | 1.91 ± 0.14bc | 1.97 ± 0.39bc | 1.46 ± 0.11ab | 1.75 ± 0.13abc | 1.71 ± 0.15abc | 2.04 ± 0.19bc |
| Ornithine | 0.98 ± 0.12ab | 2.4 ± 0.69bc | 1.74 ± 0.4abc | 2.41 ± 0.37abc | 2.67 ± 0.51c | 1.57 ± 0.23abc | 1.47 ± 0.24abc | 1.96 ± 0.39abc | 1.89 ± 0.17abc | 2.68 ± 0.43abc | 1.77 ± 0.25abc | 0.79 ± 0.11a | 1.59 ± 0.18abc | 2.19 ± 0.46abc | 5.1 ± 0.64c |
| Phenylalanine | 0.65 ± 0.22ab | 1.59 ± 0.86ab | 0.38 ± 0.05b | 0.53 ± 0.1a | 0.71 ± 0.13ab | 0.8 ± 0.12ab | 0.45 ± 0.04ab | 1.37 ± 0.74ab | 0.35 ± 0.04a | 0.6 ± 0.17ab | 0.79 ± 0.2ab | 0.48 ± 0.16ab | 0.57 ± 0.12ab | 0.7 ± 0.18ab | 1.17 ± 0.31ab |
| Phosphoric acid | 0.96 ± 0.08ab | 1.06 ± 0.05abc | 1.09 ± 0.07abc | 1.2 ± 0.05cd | 1.03 ± 0.03abc | 1.02 ± 0.07abc | 0.96 ± 0.04ab | 1.03 ± 0.07abc | 1.16 ± 0.04bcd | 1.2 ± 0.1cd | 1.04 ± 0.06abc | 0.91 ± 0.04a | 0.98 ± 0.04ab | 1.11 ± 0.07abcd | 1.29 ± 0.05d |
| Proline | 2.34 ± 0.63a | 10.55 ± 4.05abc | 18.04 ± 7.68c | 7.86 ± 3.07ab | 2.13 ± 0.55a | 3.64 ± 0.62a | 1.99 ± 0.28a | 6.92 ± 1.28ab | 18.78 ± 3.1c | 13.04 ± 1.48cd | 1.84 ± 0.36a | 1.79 ± 0.29a | 4.53 ± 0.95ab | 4.27 ± 1.06ab | 6.96 ± 1.52ab |
| Putrescine | 2.36 ± 0.83ab | 2.57 ± 0.63ab | 1.95 ± 0.12ab | 3.03 ± 0.26ab | 3.34 ± 0.5ab | 2.43 ± 0.45ab | 2.62 ± 0.5ab | 2.23 ± 0.38ab | 3.75 ± 0.45b | 2.77 ± 0.42ab | 3.58 ± 0.36ab | 1.56 ± 0.2a | 2.3 ± 0.19ab | 3.54 ± 0.29ab | 8.48 ± 2.24c |
| Pyruvate | 2.17 ± 0.22ab | 2.33 ± 0.32ab | 1.66 ± 0.21ab | 1.86 ± 0.26ab | 1.41 ± 0.08a | 2.38 ± 0.46ab | 1.87 ± 0.15ab | 2.3 ± 0.31ab | 1.93 ± 0.35ab | 1.94 ± 0.23ab | 2.78 ± 0.3b | 2.02 ± 0.18ab | 2.02 ± 0.38ab | 2.01 ± 0.33ab | 2.62 ± 0.81ab |
| Quinate | 0.76 ± 0.03bcd | 0.49 ± 0.08abc | 1 ± 0.05def | 1.03 ± 0def | 1.26 ± 0.05f | 0.82 ± 0.02cde | 0.4 ± 0.01abc | 0.39 ± 0.04abc | 1.07 ± 0.13def | 1.22 ± 0.16ef | 0.24 ± 0a | 0.52 ± 0.11abc | 0.35 ± 0.03ab | 0.34 ± 0.02ab | 1.2 ± 0.11ef |
| Rhamnose | 1.44 ± 0.23ab | 1.57 ± 0.14abc | 1.96 ± 0.14bcde | 1.9 ± 0.07abcd | 1.73 ± 0.14abc | 1.5 ± 0.23ab | 1.32 ± 0.18a | 1.51 ± 0.14ab | 2.15 ± 0.17cde | 2.35 ± 0.33de | 1.86 ± 0.19abcd | 1.35 ± 0.11ab | 1.55 ± 0.11abc | 1.73 ± 0.16abc | 2.48 ± 0.24e |
| Serine | 0.87 ± 0.25a | 1.39 ± 0.45ab | 0.6 ± 0.1a | 0.64 ± 0.13a | 0.62 ± 0.12a | 2.51 ± 0.4b | 2.49 ± 0.5b | 3.66 ± 1.01c | 0.86 ± 0.06a | 0.89 ± 0.11a | 1.36 ± 0.13ab | 1.64 ± 0.37ab | 1.26 ± 0.4a | 1.27 ± 0.32a | 1.25 ± 0.17a |
| Spermidine | 1.44 ± 0.11ab | 1.44 ± 0.28ab | 1.23 ± 0.08ab | 1.42 ± 0.22ab | 1.79 ± 0.13ab | 0.98 ± 0.15a | 1.51 ± 0.16ab | 1.19 ± 0.19a | 2.26 ± 0.33abc | 2.64 ± 0.92bc | 2.24 ± 0.29abc | 0.97 ± 0.15a | 1.31 ± 0.2ab | 2.14 ± 0.34ab | 3.54 ± 0.43c |
| Succincate | 0.55 ± 0.11a | 0.61 ± 0.06ab | 0.59 ± 0.02ab | 1.14 ± 0.4b | 0.52 ± 0.06a | 0.63 ± 0.09ab | 0.61 ± 0.12ab | 0.53 ± 0.04a | 0.74 ± 0.09ab | 0.74 ± 0.08ab | 0.53 ± 0.06a | 0.53 ± 0.06a | 0.63 ± 0.05ab | 0.53 ± 0.1a | 0.86 ± 0.17ab |
| Sucrose | 0.92 ± 0.14ab | 1.53 ± 0.28abcd | 1.11 ± 0.3abc | 1.23 ± 0.42abc | 0.81 ± 0.18a | 2.09 ± 0.2bcde | 1.66 ± 0.36abcd | 2.46 ± 0.83de | 3.04 ± 0.56e | 2.23 ± 0.51cde | 1.27 ± 0.15abcd | 1.71 ± 0.17abcd | 1.29 ± 0.22abcd | 1.65 ± 0.22abcd | 1.08 ± 0.15abc |
| Threonine | 0.57 ± 0.14ab | 1.27 ± 0.53abc | 0.41 ± 0.08a | 0.49 ± 0.06ab | 0.55 ± 0.1ab | 0.97 ± 0.15abc | 1.34 ± 0.51bc | 1.65 ± 0.49c | 0.56 ± 0.07ab | 0.89 ± 0.22abc | 0.86 ± 0.15abc | 0.66 ± 0.18ab | 0.76 ± 0.2ab | 0.78 ± 0.08ab | 1.2 ± 0.16abc |
| Trehalose | 1.89 ± 0.14ab | 2.06 ± 0.2abc | 2.05 ± 0.13abc | 2.21 ± 0.16abcd | 2.19 ± 0.12abcd | 2.14 ± 0.33abcd | 1.93 ± 0.05ab | 1.92 ± 0.25ab | 3.04 ± 0.1d | 2.87 ± 0.46cd | 2.11 ± 0.16abc | 1.76 ± 0.28a | 1.61 ± 0.11a | 1.92 ± 0.11ab | 2.82 ± 0.34bcd |
| Tryptophan | 1.69 ± 0.36abcd | 3.7 ± 1.35abcd | 2.15 ± 0.47abcd | 4.08 ± 0.71cd | 7.07 ± 0.81e | 1.54 ± 0.52abc | 0.8 ± 0.15a | 1.97 ± 0.69abcd | 2.23 ± 0.44abcd | 3.45 ± 1.64abcd | 4.02 ± 0.68bcd | 0.85 ± 0.24ab | 1.9 ± 0.41abcd | 0.96 ± 0.1abc | 4.79 ± 0.67de |
| Tyramine | 2 ± 0.44a | 1.93 ± 0.49a | 2.18 ± 0.37a | 2.14 ± 0.42a | 2 ± 0.29a | 2.69 ± 0.45ab | 2.13 ± 0.44a | 2.81 ± 0.41ab | 4.42 ± 0.47c | 3.78 ± 0.35bc | 1.69 ± 0.32a | 2.04 ± 0.31a | 5.18 ± 0.9cd | 2.41 ± 0.18ab | 6.28 ± 0.97d |
| Tyrosine | 0.61 ± 0.24ab | 0.87 ± 0.46ab | 0.32 ± 0.05a | 0.32 ± 0.03a | 0.46 ± 0.09ab | 0.79 ± 0.12ab | 0.88 ± 0.27ab | 0.39 ± 0.08ab | 0.36 ± 0.04ab | 0.41 ± 0.07ab | 0.69 ± 0.21ab | 0.29 ± 0.02a | 0.38 ± 0.09ab | 0.41 ± 0.04ab | 1.08 ± 0.21b |
| Urea | 0.76 ± 0.11a | 0.74 ± 0.1a | 0.9 ± 0.07a | 0.8 ± 0.09a | 0.7 ± 0.11a | 1 ± 0.08a | 0.85 ± 0.22a | 1.1 ± 0.11a | 1 ± 0.11a | 0.78 ± 0.15a | 0.66 ± 0.07a | 0.8 ± 0.06a | 0.92 ± 0.08a | 0.66 ± 0.11a | 0.93 ± 0.05a |
| Valine | 0.33 ± 0.12ab | 0.57 ± 0.37ab | 0.2 ± 0.04a | 0.2 ± 0.02a | 0.25 ± 0.05a | 0.87 ± 0.2bc | 0.6 ± 0.09ab | 1.17 ± 0.39c | 0.27 ± 0.04ab | 0.43 ± 0.15ab | 0.37 ± 0.05ab | 0.55 ± 0.16ab | 0.56 ± 0.2ab | 0.49 ± 0.18ab | 0.38 ± 0.06ab |
| Xylose | 2.86 ± 0.31abcd | 2.33 ± 0.35ab | 3.63 ± 0.21bcd | 5.19 ± 0.1e | 4.18 ± 0.32de | 1.76 ± 0.1a | 1.73 ± 0.46a | 1.64 ± 0.49a | 3.42 ± 0.27bcd | 3.44 ± 0.34bcd | 3.8 ± 0.45cd | 1.54 ± 0.2a | 1.93 ± 0.23a | 2.7 ± 0.57abc | 3.6 ± 0.4bcd |

**Methods S1 Growth conditions, sample collection and fruit development analysis**. The 'Ailsa Craig' tomato plants were kept under similar growing conditions as those generally described at M&M section for at least 2 months. Tomato ghost seeds were kindly provided by Dr. Marcel Kuntz and grown under similar conditions for 6-7 months, except for a lower light intensity (approx. PPFD of 100 µmol m-2 s-1) supplied in a separated zone of the same walk-in-growth chamber. Flowers were labeled at anthesis, and the date was recorded to register the number of days in fruit development (i.e. days post-anthesis, DPA). Fruits from WT and ghost plants (Fig. 1A) were harvested at the following ripening stages based on fruit size and color: mature green (MG, when reached full size but remained green, ~30 to 35 DPA), breaker (BR, ~when color change started), orange (OR) and red (R) in WT fruits; mature white (MW, ~40 DPA), breaker (BR), yellow (Y) in ghost fruits.For WT and aox1a 'MicroTom' plants, pericarp samples were harvested at the following maturity stages: Mature Green (MG), MG+1, MG+2, MG+3, MG+5, MG+7, MG+9, MG+11 and MG+13 (Fig. 4B). These stages were accurately tracked by labelling flowers on anthesis day and counting the days post-anthesis. By using this method, we accurately determined the MG stage and subsequently calculated the number of days required to reach the Breaker (BR) stage. Additionally, the transitional periods between fruit stages were visually assessed starting from the BR stage, characterized by the onset of yellow coloring on the fruit. Moreover, WT and aox1a mutant plants were monitored throughout their development and first flower buds, first anthesis flower and first fruit with a diameter of 0.3 cm were determined (Fig. 4A). Finally, fruit weight, diameter, and volume were determined in red ripe (RR) fruits as well as the total number of fruits per plant (Fig. 4A).

**Methods S2 Cloning, transformation and in vitro regeneration of CRISPR-Cas9 mutant lines.** A pair of primers for each guide was designed, denaturalized, and assembled into a pENC1.1 (pENTRY) vector previously digested with BbsI restriction enzyme. The entry vectors contained the corresponding sgRNA expression cassette flanked by Bsu36I and MluI restriction sites, and by Gateway recombinant sites to allow both types of interchange with a pDE-Cas9 plasmid (pDESTINY) providing kanamycin resistance. Agrobacterium tumefaciens GV3101 strain was used to stably transform tomato MicroTom cotyledons with plasmids harboring two sgRNAs to disrupt AOX1a genomic sequences as described previously (Fernandez et al., 2009). Primers are detailed in Supplemental Table S1. In vitro regenerated T1 lines were identified based on kanamycin resistance (100 µg ml-1) and confirmed by PCR genotyping analyses. Homozygous T2 lines lacking Cas9 were obtained after segregation.

**Methods S3 Set up of respiration and 18O discrimination analyses in fruits.** Initially, respiration analyses were performed by using liquid-phase Clark-type oxygen electrodes (Rank Brothers LTD Dual Digital Model 20). Pericarp samples from Ailsa Craig tomato fruits were cut into slices, weighed and incubated for 5, 20, and 40 min with the respiration buffer (30 mM MES, 0.2 mM CaCl2 pH 6.2) (Supplemental Fig. S1A). Thereafter, pericarp slices were placed into the oxygen electrode cuvettes containing respiration buffer and the oxygen uptake rate was measured in complete darkness at a constant temperature of 25 °C. Oxygen consumption rates were also performed after incubations with respiration buffer containing 5 mM potassium cyanide (KCN). In addition, octyl-gallate (OGAL) and salicylhydroxamic acid (SHAM) were subsequently added into the cuvette at 1mM and 20mM concentrations during measurements of KCN-inhibited tissues to compare their inhibitory effect on the AOX pathway (Supplemental Fig. S1B). All these measurements were performed before oxygen concentration reached half-air saturation levels to avoid oxygen-limiting conditions inside the cuvette (particularly important when AOX-dependent oxygen consumption is measured). Measurements of total respiration and AOX capacity in pericarp from MicroTom WT and aox1a fruits were performed similarly after approx. 15 min incubation in the respiration buffer and using 1 mM KCN (Fig. 5A). Thereafter, 18O discrimination analyses during respiration were performed by using a dual-inlet isotope ratio mass spectrometer (DI-IRMS) system as previously described (Del-Saz et al., 2017) with the following modifications for fruit tissues. After the information obtained with the oxygen electrode measurements, the sliced pericarp tissues were incubated with or without inhibitors for approx. 20 min before placing the samples in a 3 ml cuvette connected to the DI-IRMS system and an air-tight syringe containing 2 ml of air. Initially, different sizes of the sliced pericarp tissue were tested (Supplemental Fig. S2) and only those pericarp pieces of approx. or less than 2 mm thick and 1 cm length displayed no artefactual 18O discrimination by diffusion (see Del-Saz et al., 2017 for technical details related to O2 diffusion problems). To calculate the partitioning of electrons to the alternative pathway (τa), the end-point discrimination values corresponding to the AOX and COX pathways were determined in WT fruits at all developmental stages as well as in ghost fruits at the MW and Y stages (Table 1). The 18O discrimination by the AOX pathway (Δa) was determined in the presence of 5 mM KCN in all cases. In WT fruits at the MG stage, the 18O discrimination by COX was determined in the presence of 20 mM SHAM (Δc-SHAM) and it was not significantly different from the 18O discrimination in the presence of 1 mM OGAL (Δc-OGAL). Moreover, Δc-OGAL values were similar at all developmental stages tested in both WT and ghost fruits and therefore, they were used as the end-point discrimination values corresponding to the COX pathway for each developmental stage. Thereafter, the 18O discrimination in the absence of inhibitors (Δn) was determined in both WT and ghost fruits at different ripening stages (Table 1). Finally, the individual activities of the COX (νcyt) and AOX (νalt) pathways were obtained as previously described (Del-Saz et al., 2017), by multiplying the total oxygen uptake rate (Vt) and the τa (Fig.1B).

**Methods S4 RNA isolation, cDNA synthesis and RT-qPCR analyses.** RNA was isolated from lyophilized (previously frozen) pericarp tissue by using Maxwell® RSC Plant RNA Kit (Promega Biotech Ibérica, Madrid, Spain) and an automated system Maxwell® RSC Instrument (Promega Biotech Ibérica, Madrid, Spain) according to the manufacturer's instructions. RNA was quantified using a NanoDropTM 8000 spectrophotometer (Thermo Fischer Scientific) and integrity was assessed by agarose gel electrophoresis. The Prime-Script RT reagent Kit (Takara) was used to reverse transcribe 0.5 µg of extracted RNA into 20 µL of cDNA, which was subsequently diluted ten-fold and stored at -20 °C for further analysis. Relative mRNA abundance was evaluated by quantitative PCR using LightCycler 480 SYBR Green I Master Mix (Roche Basel, Switzerland) on a LightCycler 480 real-time PCR system (Roche Basel, Switzerland). Primers used and the related information are detailed in Supplemental Table S1. Two technical replicates of each biological replicate were performed, and the mean values were used for further calculations.
